# The immune landscape of murine skeletal muscle regeneration and aging

**DOI:** 10.1101/2023.11.07.565995

**Authors:** Neuza S. Sousa, Marta Bica, Margarida F. Brás, Inês B. Antunes, Isabel A. Encarnação, Tiago Costa, Inês B. Martins, Nuno L. Barbosa-Morais, Pedro Sousa-Victor, Joana Neves

## Abstract

Age-related alterations in the immune system are starting to emerge as key contributors to impairments found in aged organs. A decline in regenerative capacity is a hallmark of tissue aging, however the contribution of immune aging to regenerative failure is just starting to be explored. Here, we applied a strategy combining single-cell RNA-sequencing with flow cytometry and functional assays to perform a complete analysis of the immune environment in the aged regenerating skeletal muscle, with time and single cell resolution. Our results identified previously undescribed immune cell types in the skeletal muscle and revealed an unanticipated complexity and functional heterogeneity in immune populations, that have been regarded as homogeneous. Furthermore, we uncovered a profound remodeling of both myeloid and lymphoid compartments in aging. These discoveries challenge established notions on immune regulation of skeletal muscle regeneration, providing a new set of potential targets to improve skeletal muscle health and regenerative capacity in aging.

## INTRODUCTION

The supporting role of the immune system during regeneration is an ancient evolutionary conserved mechanism present in axolotls (*1*), insects (*2*) and teleost fish (*3-5*), which has been conserved in multiple mammalian organs (*6-11*), and is essential for regenerative success.

The adult skeletal muscle (SkM) is endowed with a remarkable regenerative capacity, serving as a paradigmatic model to understand the immune modulatory component of tissue regeneration. In the SkM, there is a temporally regulated immune response following injury, that involves immune cells of myeloid and lymphoid origin (*12*). This immune modulatory component of regeneration consists of a biphasic response involving an initial phase dominated by signals typically associated with pathogen and debris clearance, classically defined as pro-inflammatory, followed by a transition into a pro-repair phase associated with the expression of signals involved in resolution of inflammation and tissue remodeling. Immune cells have both effector functions, such as clearance of the necrotic debris, and a regulatory role over muscle stem cells (MuSCs) and other niche cells, including immune cells themselves. Effective SkM regeneration requires a proper activation of both phases of the inflammatory response, a timely transition between the two phases, and finally the resolution and cessation of inflammation in the return to tissue homeostasis (*12-14*).

Aging is a complex multifactorial process that compromises the functional integrity of tissues and simultaneously limits their regenerative potential, further contributing to the functional decline of aged organs. SkM aging manifests as a reduction in muscle mass and strength, accompanied by a striking decline in regenerative capacity, a condition broadly defined as sarcopenia (*15*). The age-related loss of SkM regenerative capacity is now viewed as a process involving both the disruption of a network of extrinsic signals regulating the function of the multiple cell types involved in the regenerative response and intrinsic alterations in MuSCs that impair their function during regeneration (*16, 17*).

The importance of the aged environment to the impairment in regenerative capacity observed in old organisms was clearly demonstrated in heterochronic parabiosis experiments, uncovering fundamental alterations in multiple circulating factors affecting regenerative capacity (*18-21*), but also pointing to a prominent role of immune cells in the modulation of these effects (*22*). Although the connection between the aging process and the immune system’s ability to maintain tissue homeostasis and repair is just beginning to be explored, recent evidence points to a central role of age-related immune alterations in the SkM’s regenerative decline (*23-26*).

To determine the full complexity of the immune response during SkM regeneration and its age-related alterations, we applied a strategy combining single-cell RNA-sequencing (scRNAseq) analysis with flow cytometry (FC) and functional assays to a time course of regenerating SkMs of young (yg) and old mice. In homeostatic SkM, we found an expansion of the lymphoid compartment in old mice, mostly attributed to the emergence of age-specific populations of B and T cells, along with an expansion of the granulocytes, dendritic cells (DCs) and specific macrophage populations. Following injury, our analysis revealed a complex landscape of macrophage activation throughout the multiple stages of SkM regeneration, including the involvement of a previously unknown population of regulatory macrophages in the early stage of SkM regeneration, and a previously unrecognized heterogeneity of macrophages associated with the reparative phase. Importantly, our analysis revealed that these new macrophage populations arise through independent differentiation trajectories, display divergent functional properties and are differentially affected by aging. Moreover, we uncovered an unanticipated diversity within the DC compartment, that was also affected by aging. Our data challenge established notions on immune regulation during SkM regeneration, revealing the participation of new cell types and states and a profound remodeling of the immune compartment in aging.

## RESULTS

### Aging is accompanied by a remodeling of the skeletal muscle immune compartment in homeostasis and during regeneration

Cells of hematopoietic origin (CD45^pos^) account for less than 5% of all mononuclear cells in homeostatic SkM, but can be expanded following injury up to 85% of all cells present during regeneration (fig. S1A). Quantification of the CD45^pos^ populations in yg and old mice, on a time course following SkM injury, confirmed a substantial increase in the CD45^pos^ population during SkM regeneration that was significantly blunted in aged mice, despite the increased number of CD45^pos^ cells in the aged homeostatic SkM (Fig. 1, A and B). Notably, the reduction in immune cells within the aged regenerating SkM was not yet detected at 1dpi, suggesting that the defects are not in the initial mechanisms of immune recruitment following injury, but rather on the subsequent processes necessary for sustaining the influx, retaining, expanding or differentiating specific immune cell types within the SkM.

**Fig. 1.**
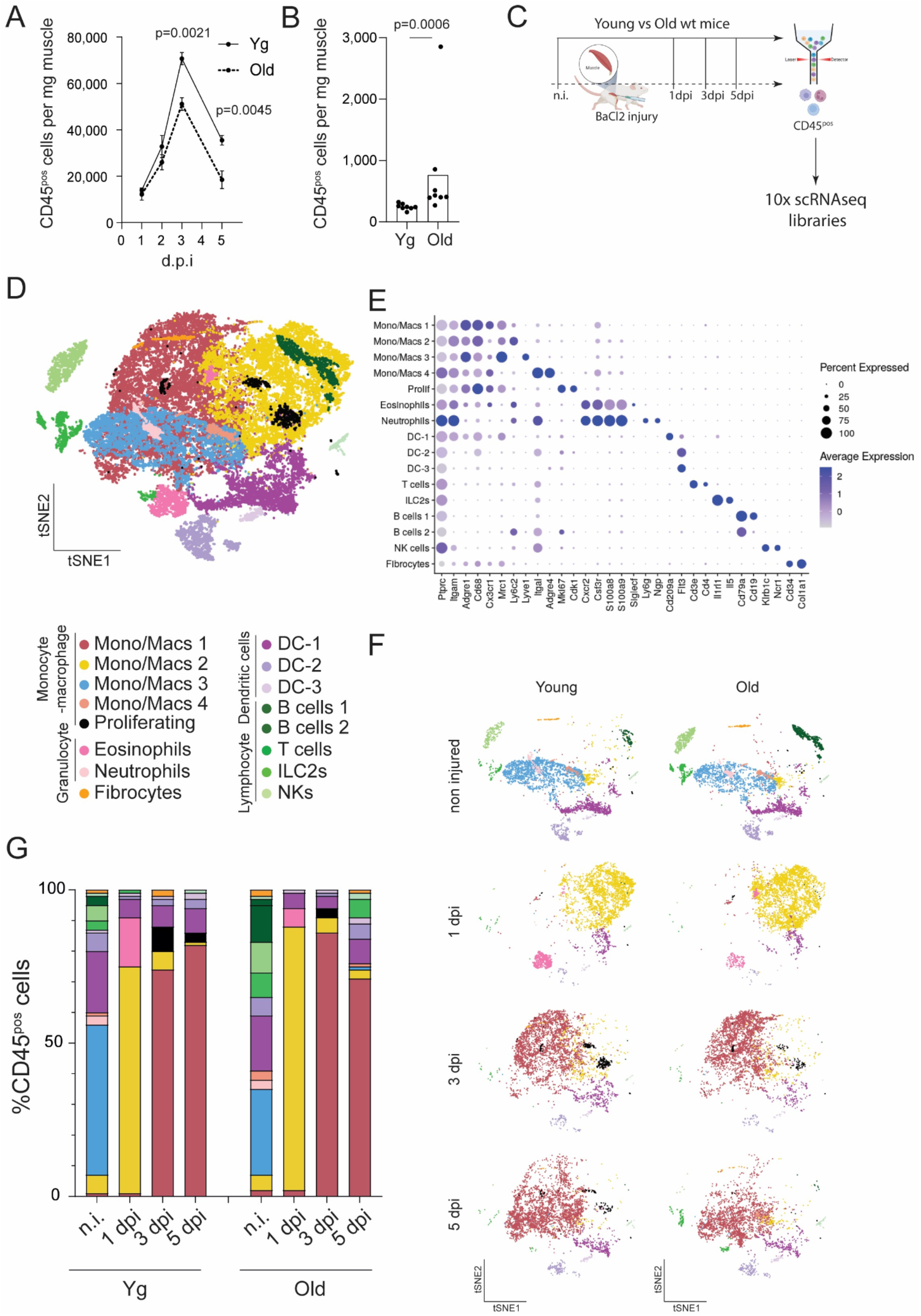
Impact of aging on the skeletal muscle’s immune compartment in homeostasis and during regeneration. **A**, Quantification, by flow cytometry, of CD45^pos^ immune cells in regenerating SkMs of yg (3– 4 months) and old (27–30 months) wt (C57BL/6) mice at different time points following injury (n=7 for yg mice at 1 dpi; n = 8 for yg mice at 2 dpi; n= 5 for yg mice at 5 dpi; n=3 for old mice at 1 dpi; n=4 for all other conditions). Data are represented as average ± s.e.m. The p values are from two-tailed Student’s t test. **B**, Quantification, by flow cytometry, of CD45^pos^ immune cells in n.i. SkMs of yg (3 months) and old (27–30 months) wt (C57BL/6) mice (n=8 for all conditions). Data are represented as average and each dot represents one animal. The p values are from two-tailed Mann-Whitney test. **C**, Workflow scheme of CD45^pos^ cell isolation, by FACS, from n.i. (n=2) and injured (1, 3 and 5 dpi; n=2/condition) SkMs of yg (3 months) and old (25 months) wt (C57BL/6) mice followed by single-cell RNA-sequencing library preparation for each sample using the 10× Chromium technology. **D**, t-distributed stochastic neighbor embedding (tSNE) visualization of unsupervised clustering (0.2 resolution) of the CD45^pos^ populations isolated from SkMs from all eight conditions (n.i., 1, 3 and 5 dpi of yg and old wt (C57BL/6) mice). Colors correspond to clustering of unique immune cell populations. **E**, Dotplot showing scaled average gene expression (color) and percentage of cells (size) expressing featured markers of each of the sixteen clusters represented in D. **F**, tSNE visualization of unsupervised clustering (0.2 resolution) of the CD45^pos^ populations isolated from SkMs of yg and old mice, as shown in D, separated by time point and age. Color codes as shown in D. **G**, Relative proportions of the different cell types identified by the scRNAseq analysis in n.i. and injured (1, 3 and 5 dpi) SkMs of yg and old mice. Color codes as shown in D. SkM, skeletal muscle; yg, young; wt, wild-type; dpi, days post injury; n.i., non-injured; FACS, Fluorescence-activated Cell Sorting; Mono, monocyte; Macs, macrophage; DCs, dendritic cells; ILC2s, innate lymphoid cells type 2; NKs, natural killer cells.

To characterize the impact of aging on the diversity of the immune environment in the SkM, and on the dynamics of the immune response associated with SkM regeneration, we fluorescence-activated cell sorting (FACS)-purified the CD45^pos^ population from non-injured (n.i.) and injured SkM at 1, 3 and 5 days post-injury (dpi) from yg (3mo) and aged (25mo) mice and prepared scRNAseq libraries for each sample using 10× Chromium technology (Fig. 1C). Sequencing metrics and quality control measures for each library are presented in the Supplementary Materials (fig. S1B-E).

Using a total of 40,956 individual cells derived from all 8 conditions, we performed unsupervised graph-based clustering, using the top 30 principal components (PCs), and projected them onto a t-distributed stochastic neighbor embedding (tSNE) plot. An initial analysis grouped our cells into sixteen unique clusters (Fig. 1D). Among the identified clusters, half of them expressed the common myeloid marker *Itgam* (also known as CD11b) and corresponded to distinct populations of myeloid cells, identified based on hallmark gene expression (Fig. 1E). The three biggest clusters represented cells belonging to the monocyte/macrophage lineage (Mono/Macs 1-3), identified based on the expression of their common markers: *Adre1* (also known as F4/80), *Cd68* and *Mrc1* (also known as CD206) (Fig. 1, D and E and fig. S2A). The Mono/Mac clusters 1 and 2 expressed *Ly6c2* and were mostly present in injured SkM (Fig. 1, D-F and fig. S2A), corresponding to infiltrating monocyte-derived macrophages that dominate the immune response during SkM regeneration. Mono-Mac cluster 1 was present at later stages of SkM regeneration (3 and 5 dpi), while Mono-Mac cluster 2 was mostly detected at 1 dpi (Fig. 1F), suggesting that they represent the broadly defined populations of pro-repair and pro-inflammatory macrophages, respectively. Indeed, Mono/Macs 1 and 2 were distinguished by the expression levels of *Cx3cr1* and *Ccr2* (fig. S2B), and *Ly6c2* expression was higher in Mono/Macs 2, in agreement with the expression profile of these populations (*14*). Mono/Mac cluster 3 expressed the *Lyve1* marker and was mostly represented in n.i. SkM (Fig. 1, D-F and fig. S2A), corresponding to a recently described population of SkM resident macrophages (*27*). The Mono/Mac compartment also included two smaller clusters: the Mono/Mac cluster 4 and a cluster of proliferating cells (Fig. 1D). The latter was enriched in genes associated with cell cycle progression, *Mki67* and *Cdk1*, and was mostly present in regenerating SkMs (Fig. 1, D-F), likely corresponding to proliferating macrophages (fig. S2A). The Mono/Mac cluster 4 was mostly present in n.i. SkM and was enriched in specific markers, *Itgal* (also known as CD11a) and *Adre4* (Fig. 1, D-F), corresponding to an yet unidentified population of SkM resident cells of the monocyte/macrophage lineage.

Among the myeloid cells, we also identified 2 clusters of granulocytes, expressing the lineage-specific markers *Cxcr2*, *Csf3r*, *S100a8* and *S100a9* and corresponding to two different cell types, neutrophils and eosinophils, distinguished by the expression of *Ngp* and *Ly6*g, or *Siglecf*, respectively (Fig. 1, D and E and fig. S2A). Our analysis also identified three clusters of cells belonging to the dendritic cell (DC) lineage (DC-1, DC-2 and DC-3), defined based on the expression of the common DC markers, and previously uncharacterized in the SkM. Cluster DC-1, characterized by enriched *Cd209a* and *Itgam* expression, was clearly differentiated from clusters DC-2 and 3, which exhibited elevated *Flt3* levels but lacked myeloid lineage genes. (Fig. 1, D and E). Notably, our analysis revealed that the genes commonly used to identify monocyte and macrophage subpopulations in the SkM are expressed by other myeloid cell populations (fig. S2A), including DCs and eosinophils, highlighting the need to establish new markers that define macrophage populations with higher specificity.

Among the CD11b^neg^ cells present in our dataset, we identified five different clusters of cells belonging to the lymphoid compartment and one population of immune cells previously uncharacterized in the SkM, identified as fibrocytes. The lymphoid cell types were mostly represented in homeostasis and included one cluster of T cells, expressing *Cd3e* and *Cd4*, two clusters of B cells expressing *Cd79a* and *Cd19,* one cluster of Natural Killer cells (NKs), expressing *Klrb1c* (also known as Nkx1.1) and *Ncr1*, and one cluster of Innate lymphoid cells type 2, expressing *Il1rl1*(also known as ST2) and *Il5* (Fig. 1, D-F). Fibrocytes are cells of hematopoietic origin (CD45^pos^) that co-express markers of the mesenchymal lineage (*28*), and were identified in our dataset based on the expression of *Cd34* and *Col1a1* (Fig. 1, D and E). Decomposition of our full scRNAseq dataset by time point and age revealed a profound remodeling of the immune compartment following SkM injury and suggested previously unknown effects of aging on immune population dynamics and states within both the homeostatic and regenerating SkM (Fig. 1F and G). In yg mice, the CD45^pos^ population present in n.i. SkM was primarily composed of resident macrophages (Mono/Macs 3, 49%), while the remaining half consisted of infiltrating macrophages (Mono/Macs 2, 6%), Neutrophils (3%), DCs 1-3 (27%), T cells (3%), ILC2s (5%), B cells (3%), NKs (1%) and Fibrocytes (1%) (Fig. 1, F and G). Immediately after injury (1 dpi), the SkM immune compartment was dominated by infiltrating macrophages (Mono/Macs 2, 74%), eosinophils (16%) and DCs 1-3 (8%) (Fig. 1, F and G). By 3 dpi, the immune population was almost entirely composed of Mono/Macs 1 (74%), and only 6% of the Mono/Mac-2 subpopulation remained, together with 8% of proliferating macrophages and 10% of DCs 1-3 (Fig. 1, F and G). Finally, at 5 dpi the population of proliferating macrophages was reduced to 3% and the infiltrating population of Mono/Macs-2 to 1%, with Mono/Macs1 representing 82% and DCs 1-3 13% of the CD45^pos^ cells (Fig. 1, F and G).

The aged homeostatic SkM displayed a shift in the balance between myeloid and lymphoid cell types, with a 3-fold expansion of the lymphoid compartment and a relative contraction of the resident macrophage population. Moreover, we observed a 2-fold increase in the representation of Fibrocytes and Mono/Mac cluster 4 in aged n.i. SkM (Fig. 1, F and G). Notable age-related alterations during SkM regeneration included a 3-fold reduction of eosinophil representation at 1dpi, a 3-fold reduction in the proliferating macrophage population at 3 dpi and a 6-fold expansion of the lymphoid compartment at 5dpi (Fig. 1, F and G), potentially recapitulating the changes in homeostatic conditions.

This initial analysis revealed important age-related changes in the immune milieu of the SkM in homeostasis and regeneration, which may be relevant for the deterioration of SkM health and regenerative capacity in aging. To understand the dynamics of these different immune cell types during regeneration and aging, we zoomed in on individual classes of immune cells, exploring our data with increased detail.

### The lymphoid compartment is expanded in the aged homeostatic skeletal muscle

Lymphoid cells represented only 12% of the immune compartment in yg homeostatic SkM but were expanded in old animals, where they corresponded to approximately one third of the CD45^pos^ cells (Fig. 1G and Fig. 2A). All lymphoid cell types, except for the NKs, were overrepresented in aged SkMs (Fig. 1, F and G and Fig. 2A). Aging also altered the relative proportions of lymphoid cells, resulting in B cells becoming the predominant lymphoid cell type in aged SkMs (Fig. 2B).

**Fig. 2.**
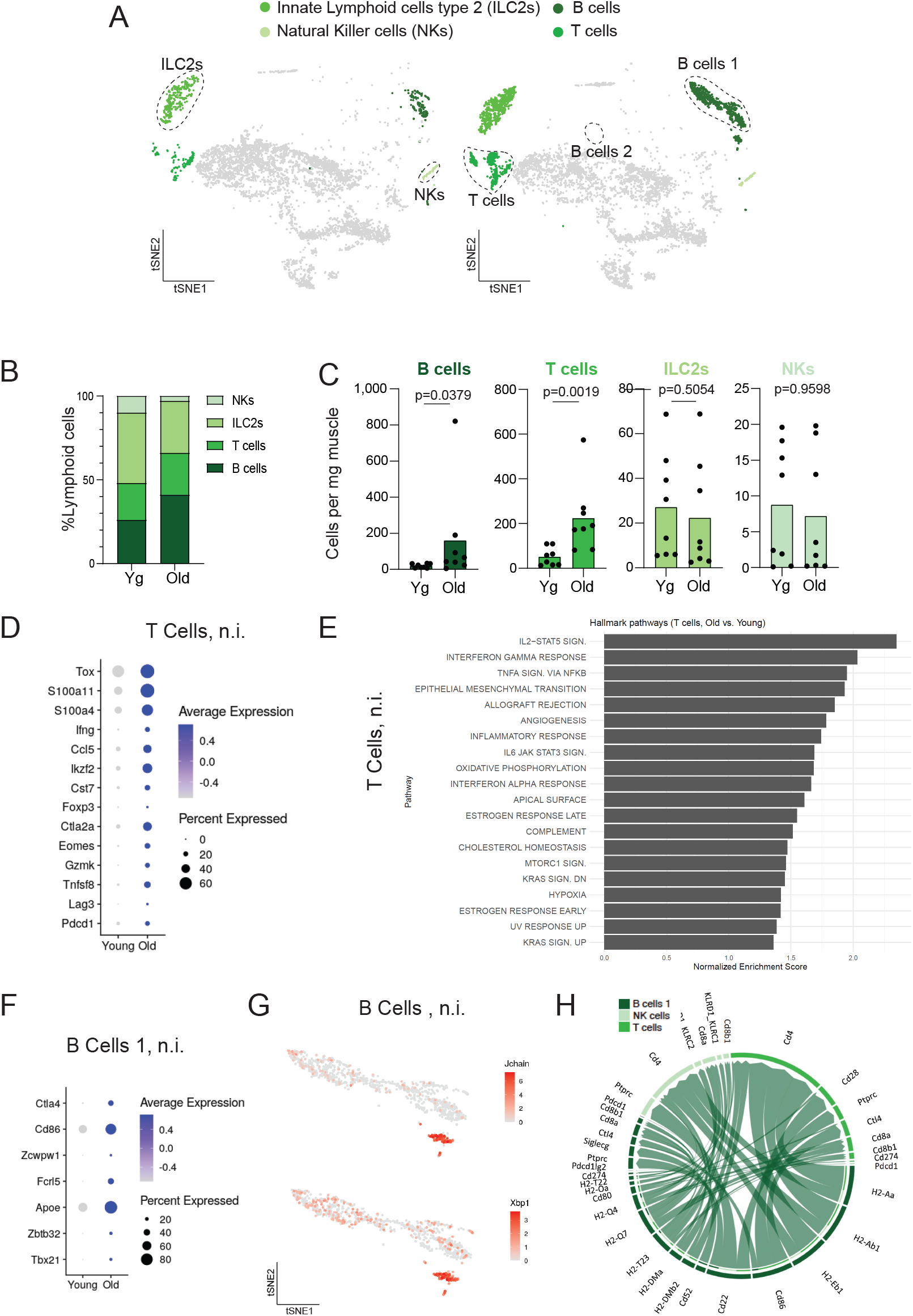
Impact of aging on the lymphoid compartment in homeostatic skeletal muscle. **A,** tSNE plots (0.2 resolution) displaying the lymphoid populations identified in n.i. SkMs of yg (3 months) and old (25 months) wt (C57BL/6) mice. Each colour represents one lymphoid population as described in the legend. Non-lymphoid populations are shown in grey. **B,** Relative proportions of lymphoid populations (NKs, ILC2s, T and B cells) identified by the scRNAseq analysis, in n.i. SkMs of yg and old wt mice. **C,** Quantification, by flow cytometry, of B cells, T cells, ILC2s and NKs in n.i. SkMs of yg (3 months) and old (27-30 months) wt (C57BL/6) mice (n=8 for all conditions). Data are represented as average, and each dot represents one animal. The p values are from two-tailed Mann-Whitney test. **D,** Dotplot showing scaled average gene expression (color) and the percentage of T cells (size) expressing hallmark genes associated with T cell aging, in n.i. SkMs of yg and old wt mice. **E,** Normalized enrichment score (NES) of MSigDB hallmark pathways between yg (control) and old T cells in n.i. SkMs of wt mice. The pathways in display were selected for NES > 0 and adjusted p-value < 0.05. See also Fig. S3E. **F,** Dotplot showing scaled average gene expression (color) and the percentage of B cells (size) expressing hallmark genes associated with T cell aging, in n.i. SkMs of yg and old wt mice. **G,** tSNE plot showing the normalized and log-transformed expression levels of *Jchain* and *Xbp1* genes in the B cell compartment of n.i. SkMs of aged mice. **H,** CellChat chord diagram illustrating up-regulated ligand-receptor interactions between B cells and other lymphoid cell types based on the differentially expressed genes of each cluster with aging in n.i. SkMs. Gene colour represents the cell type where they are differentially expressed, and the arrows in the diagram point from a given ligand to its associated receptor. Arrow width is proportional to the communication probabilities computed by CellChat for the interactions shown in the diagram. SkM, skeletal muscle; yg, young; wt, wild-type; n.i., non-injured; ILC2s, innate lymphoid cells type 2; NKs, natural killer cells.

To validate these findings and understand whether the overrepresentation of the lymphoid compartment in our scRNAseq dataset was due to a loss of myeloid cells, or there was indeed an expansion of specific lymphoid cell types, we quantified, by FC, T cells (CD45^pos^CD3^pos^), B cells (CD45^pos^CD3^neg^CD79^pos^), ILC2s (CD45^pos^CD3^neg^ST2(*Il1rl1*)^pos^) and NKs (CD45^pos^Nk1.1^pos^) in n.i. SkMs of yg and aged mice (fig. S3, A-C). Our results showed that B cells and T cells significantly increased in number in old SkMs while ILC2 and NK abundance remained unaltered (Fig. 2C), explaining the remodeling of the lymphoid compartment predicted by our scRNAseq data (Fig. 2B).

To understand the age-related increase in T and B cell numbers in the aged SkM, we used the scRNAseq data to discriminate the identity of the cells contributing to their expansion. Within the T cell compartment, CD4^pos^ T cells were the predominant subtype, but CD8^pos^ T cells, Tregs (Foxp3^pos^) and γδTcells (TCRg-C1^pos^) were also detected, and all subpopulations were expanded in aged SkMs (fig. S3D). The expansion of the T cell population was associated with the appearance of three age-associated CD4^pos^ subsets, identified in other aged organs (*29, 30*): exhausted (*Tox*, *Pdcd1, Lag3, Tnfsf8)*, cytotoxic (*Gzmk*, *Eomes, Ctla2a*) and activated Tregs (*Foxp3, Tnsrsf9, Cst7, Ikzf2*) (Fig. 2D). In addition, our aged cluster was also enriched in a PD1^pos^GZMK^pos^ subset of exhausted CD8^pos^ T cells (Fig. 2D, (*31*)). These T cell subsets were further enriched in proinflammatory molecules (*Ccl5, Ifng, S100a4, S100a11*), a feature shared with T cells from other aged organs (Fig. 2D, (*29, 30*)), suggesting a potential contribution of T cells to the established inflammatory environment of the aged SkM. Consistently, gene set enrichment analysis of the dataset of genes upregulated in aged T cells revealed IL2/STAT5 signaling pathway as most activated by aging (Fig. 2E and fig. S3E). This pathway has been associated with the increased propensity of aged CD4^pos^ T cells to initiate effector programs (*32*).

The B cell compartment in yg SkMs was mostly composed by naïve B cells (*Fcer2a, Satb1*, Fig. 2A and fig. S3F). The expansion of B cells in aging could be mostly attributed to the emergence of a new population (*Tbx21, Zbtb32, Apoe, Fcrl5,* Fig. 2, A and F and fig. S3F), distinct from the conventional naïve and memory subtypes, but also found in other aged tissues and termed “age-associated B cells” (*29, 31, 33*). Notably, we also found an increase in expression of *Zcwpw1*, *Cd86* and *Ctla4* (Fig. 2F), which can be attributed to a subset of peritoneal B1-like cells which become activated with age and serve as inducers of inflammatory phenotypes in cytotoxic CD8^pos^ T cells (*29, 31, 34*). In addition, the aged B cell compartment also included a cluster of cells expressing the markers of plasma B cells (*Jchain, Xbp1,Derl3*, Fig. 2G), also found in the spleen, bone-marrow, kidney and adipose tissue of aged mice (*29, 35*). To understand the consequences of the age-related alterations in B cells on the SkM immune environment, we performed an analysis of intercellular communication using CellChat (*36*). With aging, we found a maximum of differential number of interactions originating in B cells affecting other B cells and a maximal interaction strength towards T cells and NK cells (fig. S3G). Interestingly, these altered interactions correspond mostly to an increase in cell-cell contact signaling medicated by Major Histocompatibility Complex II (MHCII, Fig. 2H).

Due to the high influx of myeloid cells during SkM regeneration, our scRNAseq data did not allow us to detect enough lymphoid cells at 1, 3 and 5 dpi to extract conclusions regarding the dynamics of the different cell types during regeneration or their potential age-related impairments. To overcome this limitation, we quantified the dynamic change of T and B cells during regeneration by FC in yg and old mice. T cell number progressively increased after injury peaking at 3dpi, in agreement with previous data (*37, 38*) and this dynamic was not changed in aged animals (Fig. S3H), while B cell numbers increased mostly between 1 and 2 dpi but failed to accumulate in aged SkMs (Fig. S3I).

These results revealed an unanticipated alteration of the lymphoid compartment in the aged SkM that has the potential to impact SkM inflammatory status, and possibly affect SkM health.

### Aging blunts the DC response during skeletal muscle regeneration

Dendritic cells (DCs) represented about one quarter of the CD45^pos^ population in homeostatic SkM in both yg and old animals. During regeneration, the contribution of the DC population to the immune compartment was reduced to about 5% at 1 dpi and progressively increased during the regenerative process, without major age-related changes in these dynamics (Fig. 1G and fig. S4A).

Analysis of the DC population with increased resolution allowed the discrimination of four DC subtypes based on hallmark gene expression (Fig. 3, A and B and fig. S4B). The majority of DCs present in the SkM expressed the Dendritic Cell-Specific Intercellular adhesion molecule-3-Grabbing Non-integrin (DC-SIGN, also known as CD209) and represented 2 distinct sub-populations: monocyte-derived dendritic cells (moDCs) and plasmacytoid dendritic cells (pDCs), discriminated by the expression of *Mrc1*(also known as *Cd206*) and *Dcstamp* or *Ly6c* and *Cd7*, respectively (Fig. 3, A and B and fig. S4B) (*39-43*). Interestingly, moDCs were mostly detected in homeostatic SkM, while pDCs became the most abundant DC subtype during regeneration (Fig. 3C and fig. S4C), representing in each case approximately 60-80% of the DC population. In addition, two other DC subtypes could be detected in homoeostatic and regenerating SkM that were CD209^neg^, including conventional DCs type 1 (cDC1s), expressing *Itgae* (also known as *Cd103*), *Xcr1* and *Clec9a* (*39*), and a minor subset identified based on the expression of *Ccr7*, *Fscn1*, and *Mreg*, resembling a recently described population of mature DCs enriched in immunoregulatory molecules (mregDCs) (*44*) (Fig. 3, A and B and fig. S4B). cDC1s represented about 20-30% of the DC population, were reduced relative to others at 1dpi, and then progressively enriched during regeneration (Fig. 3C and fig. S4C). mregDCs represented less than 10% of the DC population in homeostasis and during the early stages of SkM regeneration, and were expanded at 5 dpi (Fig. 3C and fig. S4C).

**Fig. 3.**
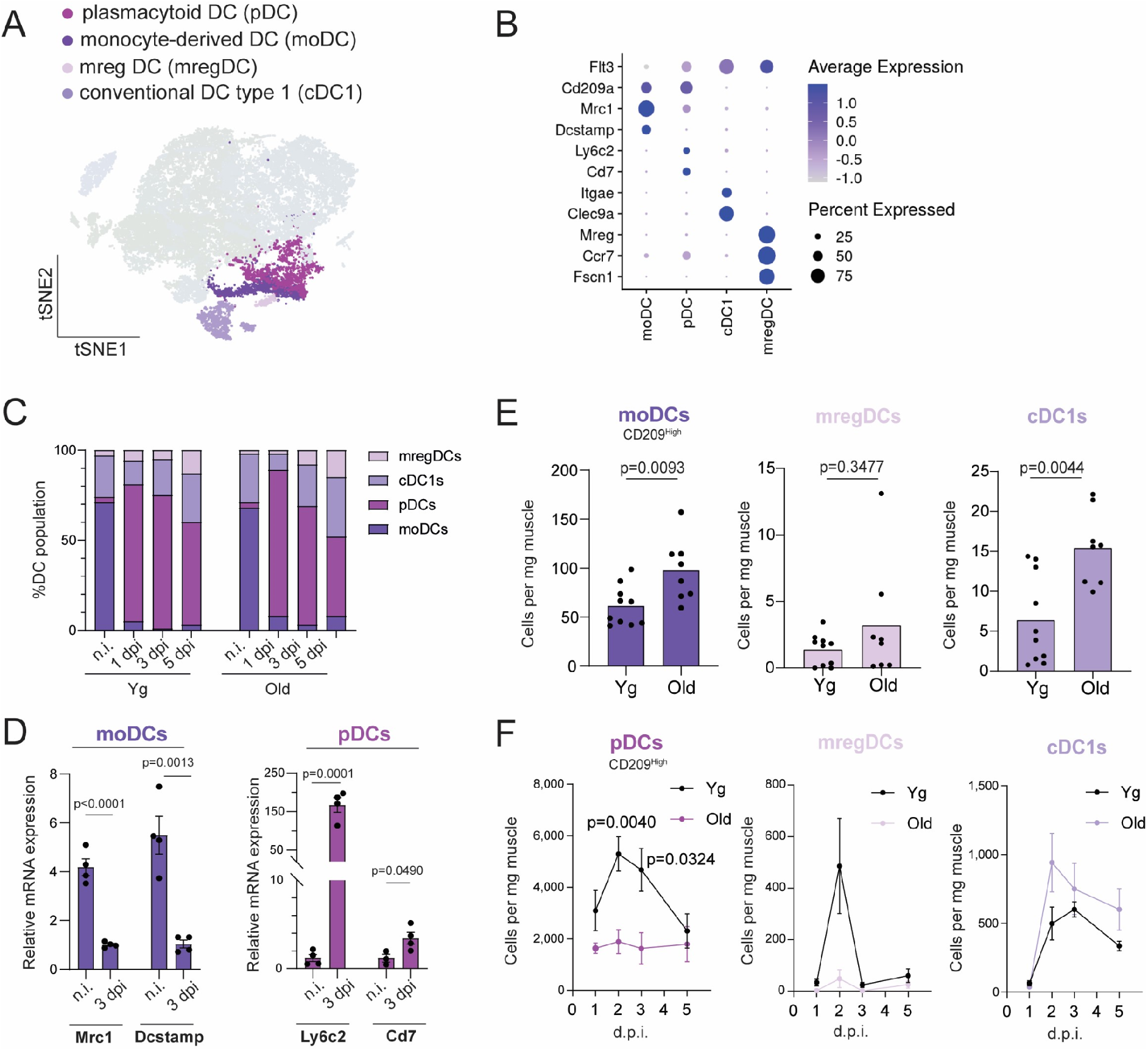
Impact of aging on skeletal musclés dendritic cell populations in homeostasis and during regeneration. **A**, tSNE plots (0.4 resolution) displaying the four unique clusters of DC populations identified in SkMs from all eight conditions (n.i., 1, 3 and 5 dpi, of yg and old wt (C57BL/6) mice). Colors represent clusters of subtypes of DCs. Non-DC clusters are shown in grey. **B**, Dotplot showing scaled average gene expression (color) and percentage of cells (size) expressing featured markers of each of the four DCs subtypes represented in A. **C**, Relative proportions of DC subtypes identified by the scRNAseq analysis, in n.i. and injured (1, 3 and 5 dpi) SkMs of yg and old mice. **D**, Relative levels of Mrc1, Dcstamp, Ly6c2 and Cd7 mRNA, detected by RT-qPCR, in CD209^high^ DCs isolated, by FACS, from n.i and 3 dpi SkMs of yg (3 months) wt (C57BL/6) mice (n=3 for n.i of Cd7 analysis; n=4 for all others analysis). **E,** Quantification, by flow cytometry, of moDCd, mregDCs and cDC1s, in n.i. SkMs of yg (3 months) and old (27–30 months) wt (C57BL/6) mice (n=10 for yg mice and n=8 for old mice). **F**, Quantification, by flow cytometry, of pDCs, mregDCs and cDC1s, in regenerating SkMs of yg (3-4 months) and old (27–30 months) wt (C57BL/6) mice at different time points following injury (n=3 for old mice at 1 dpi; n=8 for yg mice at 2 dpi; n=7 for yg mice at 3 dpi; n=4 for old mice at 2 and 3 dpi; n=5 for all other conditions). In D and F, data are represented as average ± s.e.m, and in D each dot represents one animal. In E, data are represented as average, and each dot represents one animal. In D, p-values are from two-tailed Student’s t test. In E, p values in moDCs analysis are from two-tailed Student’s t test, p value in mregDCs and cDC1s analysis is from two-tailed Mann-Whitney test. In F, p value of 2dpi analysis is from two-tailed Mann-Whitney test, and p value of 3dpi analysis is from two-tailed Student’s t test. SkM, skeletal muscle; yg, young; wt, wild-type; n.i., non-injured; dpi, days post injury; DCs, dendritic cells; moDCs, monocyte-derived dendritic cells; pDCs, plasmacytoid dendritic cells; cDC1s, conventional dendritic cells type 1; mregDCs, mature dendritic cells enriched in immunoregulatory molecules.

To evaluate the dynamics of the DC populations during SkM regeneration in yg and old animals and validate the findings from our scRNAseq analysis, we designed a FC panel using cell surface markers based on the hallmark gene expression described above, including CD209, CD103, XCR1 and CCR7 (fig. S4D). Since pDCs and moDCs were defined by the expression of the same surface marker (CD209) and are almost exclusively present in n.i. and injured SkMs, respectively, we performed RT-qPCR analysis on the sorted fraction of both conditions to validate their identity. Indeed, CD209^high^ cells sorted from n.i. SkMs were enriched in *Mrc1* and *Dcstamp*, while CD209^high^ cells sorted from SkM at 3dpi were enriched in *Ly6c2* and *Cd7* (Fig. 3D). Our FC analysis showed that in n.i. SkMs the numbers of moDCs and cDC1s were significantly increased with age (Fig. 3E). Moreover, aged SkMs displayed significant defects in the repair-associated DC response (Fig. 3F): while in yg SkMs, pDCs, mregDCs and cDC1s were dynamically expanded after injury, there was a striking defect in the pDC response in aged animals. These data revealed an underappreciated complexity of the DC compartment in the SkM and important age-related defects in the DC dynamics during regeneration with potential consequences for SkM regeneration yet unexplored.

### Granulocytes have an altered profile and display severe recruitment defects during regeneration in the aged skeletal muscle

Two distinct populations of granulocytes could be identified in our scRNAseq dataset, represented only in the n.i. and 1 dpi SkM: Neutrophils (Ly6G^pos^) and Eosinophils (SiglecF^pos^) (Fig. 1, D and E and Fig. 4A). Granulocytes are difficult to capture through scRNAseq technologies due to their low RNA content and propensity to undergo apoptosis shortly after isolation, which may explain why we were unable to detect neutrophils in 1 dpi SkMs, when they are known to be a highly abundant cell type (*12*). To overcome this limitation and understand the impact of aging on the dynamics of granulocyte populations, we used FC to identify Neutrophils (Cd11b^pos^Ly6G^pos^) and Eosinophils (Cd11b^pos^SiglecF^pos^) in the SkM of yg and aged mice (fig. S5A).

**Fig. 4.**
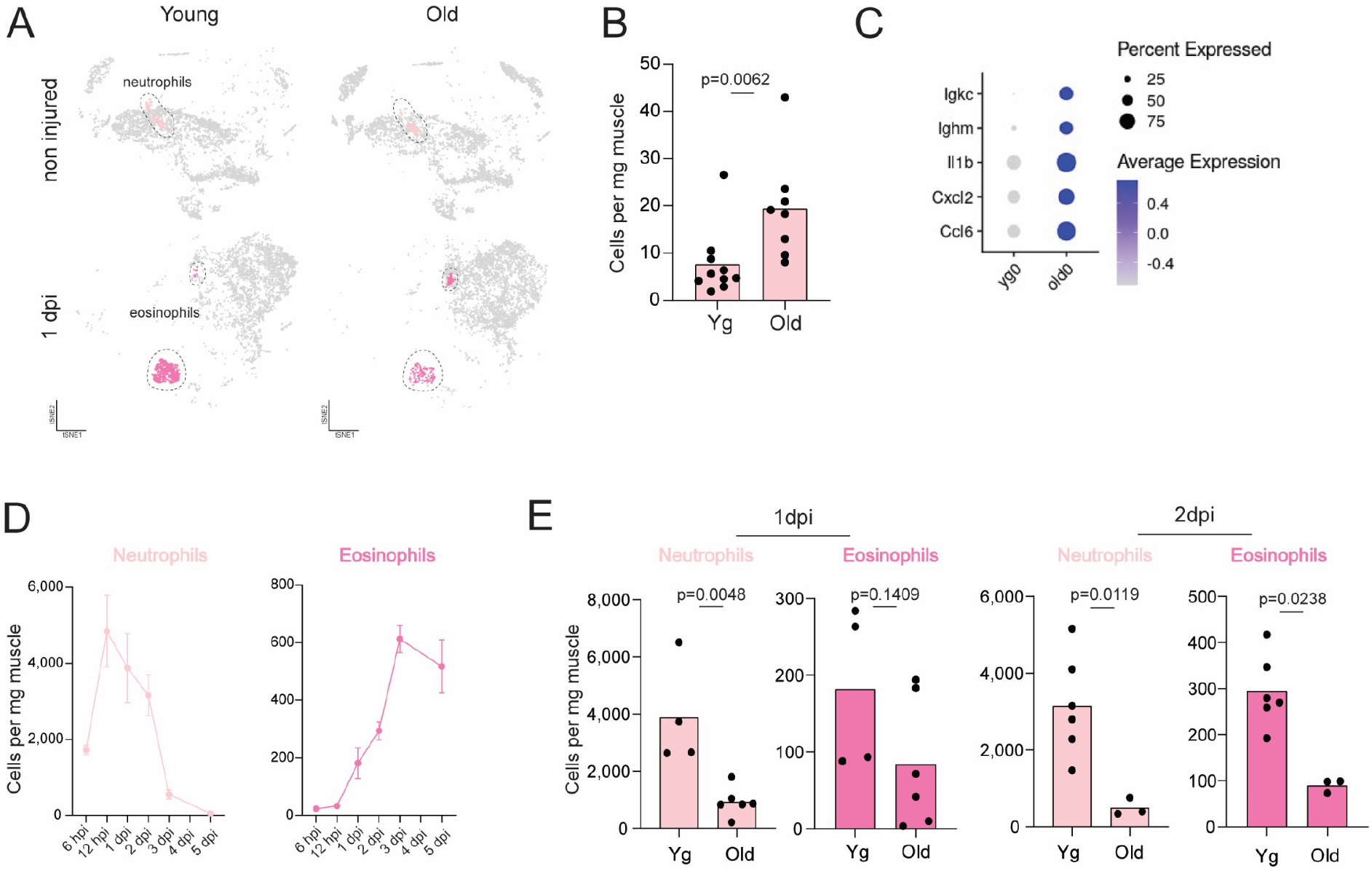
Impact of aging on skeletal muscle’s neutrophils and eosinophils populations in homeostasis and during regeneration. **A**, tSNE plots (0.2 resolution) displaying the neutrophils and eosinophils populations identified in SkMs from n.i. and 1 dpi SkMs of yg and old mice wt (C57BL/6) mice. Colors represent clusters of neutrophils and eosinophils. Non-granulocyte clusters are shown in grey. **B,** Quantification, by flow cytometry, of neutrophils in n.i. SkMs of yg (3 months) and old (27-30 months) wt (C57BL/6) mice (n=10 for yg mice and n=8 for old mice). **C,** Dotplot showing scaled average gene expression (color) and the percentage of neutrophils (size) expressing hallmark genes associated with neutrophil aging, in n.i. SkMs of yg and old wt mice. **D,** Quantification, by flow cytometry, of neutrophils and eosinophils in regenerating SkMs of yg (2–4 months) wt (C57BL/6) mice at different timepoints following injury (n=6 for 2dpi, n=10 for 3dpi and n=4 for all other conditions). **E,** Quantification, by flow cytometry, of neutrophils and eosinophils in regenerating SkMs of yg (3–4 months) and old (26-29 months) wt (C57BL/6) mice, at 1dpi and 2dpi (n=4 for yg mice at 1dpi; n=3 for old mice at 2dpi, n=6 for all other conditions). In D, data are represented as average ± s.e.m. In B and E, data are represented as average and each dot represents one animal. In B, p-values are from two-tailed Mann-Whitney test. In E, p values in neutrophils and eosinophils anal–ysis at 1 dpi and neutrophils at 2 dpi are from two-tailed Student’s t test, and the p value in eosinophils analysis at 2 dpi are from two-tailed Mann-Whitney test. SkM, skeletal muscle; yg, young; wt, wild-type; n.i., non-injured; dpi, days post injury; hpi, hours post injury.

In homeostatic SkM, only neutrophils were detected in our scRNAseq dataset and our FC analysis revealed an expansion of this neutrophil population in aged mice (Fig. 4B), in agreement with previous reports (*24*). Importantly, aged neutrophils also displayed aberrant expression of immunoglobulins, along with up-regulation of genes associated with chemotaxis and inflammation (Fig. 4C).

The expansion of neutrophils in the homeostatic SkM of aged mice contrasted with the severe defects in granulocyte recruitment after injury. Indeed, our FC analysis in yg mice, revealed an immediate wave of neutrophil recruitment peaking at 1 dpi, followed by the recruitment of eosinophils starting at 1dpi with a maximum accumulation at 3dpi (Fig. 4D). Notably, the number of neutrophils and eosinophils was significantly lower in aged SkM in the early time points following injury (Fig. 4E), suggesting defects in the mechanism that attract or retain these cell populations in the regenerating SkM. The striking reduction in eosinophils observed in our FC analysis validates the results obtained in our scRNAseq analysis (Fig. 1G and Fig. 4A), revealing a novel important defect in the repair-associated myeloid response to be explored in the context of SkM regenerative failure in aging.

### A complex landscape of macrophage states participates in skeletal muscle regeneration and is disrupted in aged animals

Cells belonging to the monocyte/macrophage lineage were the most represented cell types in our scRNAseq dataset (∼60%), allowing for a deeper characterization of their cellular states. Re-clustering of Mono/Macs 1-4 populations and analysis at higher resolution distinguished multiple macrophage states associated with different phases of the regenerative process (Fig. 5, A-C).

**Fig. 5.**
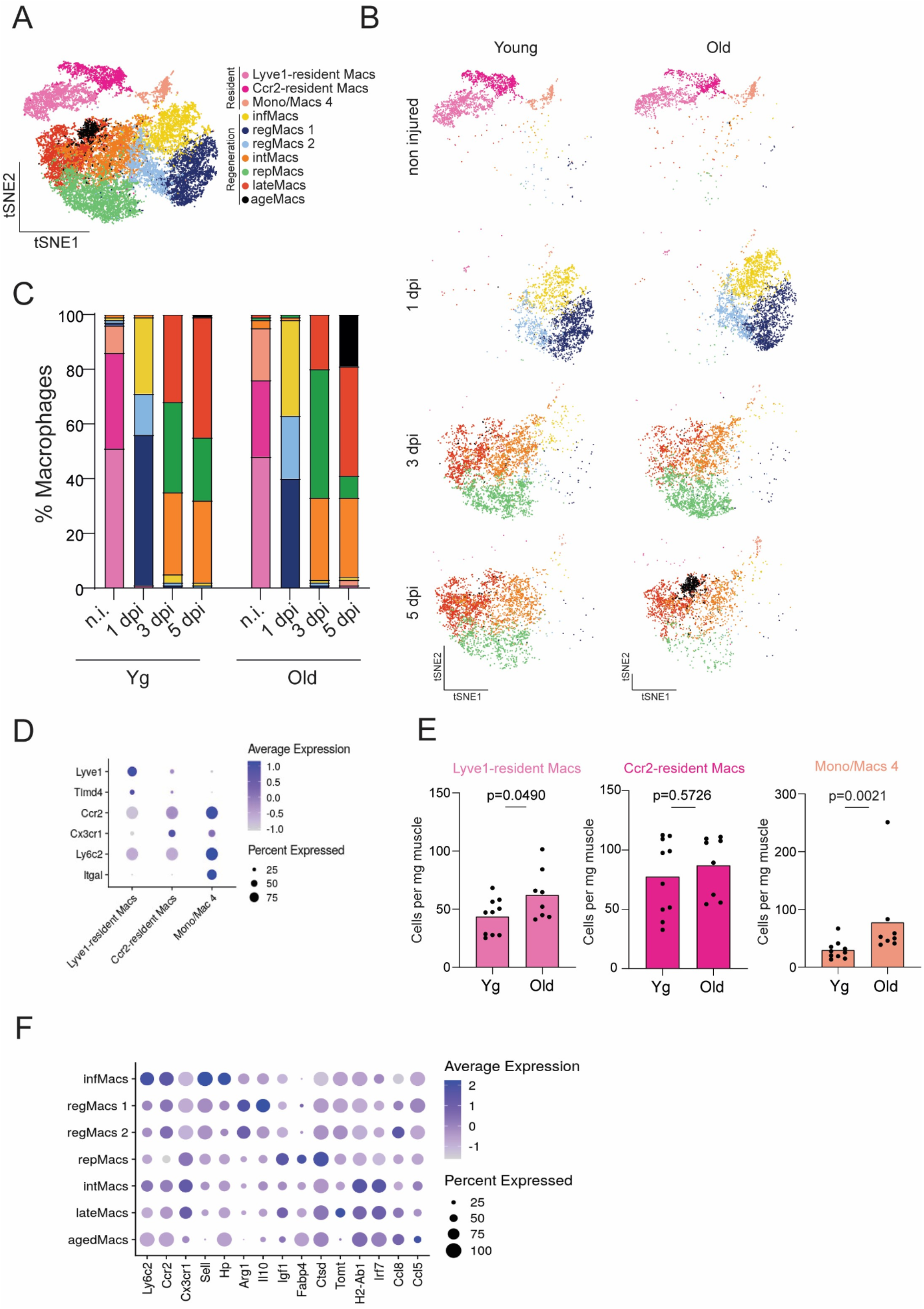
Impact of aging on skeletal muscle’s macrophage diversity and dynamics in homeostasis and during regeneration. **A,** t-distributed stochastic neighbor embedding (tSNE) visualization of unsupervised re-clustering (0.6 resolution) of Mono/Macs1-4 populations identified in Fig. 1D, from all eight conditions (n.i, 1, 3 and 5 dpi of yg (3 months) and old (25 months) wt (C57BL/6) mice). Colors correspond to clustering of unique macrophage populations. **B,** tSNE visualization of unsupervised re-clustering (0.6 resolution) of Mono/Macs1-4 populations, as shown in A, separated by time point and age. Color codes as shown in A. **C,** Relative proportions of the different macrophage populations identified by the scRNAseq analysis in n.i. and injured (1, 3 and 5 dpi) SkMs of yg and old mice. Color codes as shown in A. **D,** Dotplot showing scaled average gene expression (color) and percentage of cells (size) expressing featured markers of each of the three macrophage populations present in n.i. SkMs of yg and old mice, as shown in B. **E,** Quantification, by flow cytometry, of resident macrophage populations and Mono/macs 4, in n.i. SkMs of yg (3 months) and old (27–30 months) wt (C57BL/6) mice (n=10 for yg mice and n=8 for old mice). Data are represented as average, and each dot represents one animal. For Lyve1-resident Macs, p value is from two-tailed Student’s t test. For Ccr2-resident Macs and Mono/Macs 4, p-value is from two-tailed Mann-Whitney test. **F,** Dotplot showing scaled average gene expression (color) and percentage of cells (size) expressing featured markers of each of the seven macrophage populations present in injured (1, 3 and 5 dpi) SkMs of yg and old mice, as shown in B. SkM, skeletal muscle; yg, young; wt, wild-type; n.i., non-injured; dpi, days post injury; Mono, monocyte; Macs, macrophages.

In n.i. SkM, our re-clustering analysis discriminated three populations, confirming and expanding recently published studies (*27, 45, 46*). These clusters resulted from the discrimination of two populations of Ly6C^neg^ resident macrophages, distinguished by the expression of *Lyve1* and *Timd4* (Lyve1-resident Macs) or *Ccr2* and *Cx3cr1* (Ccr2-resident Macs) and the cluster of Mono/Mac 4, which expressed high levels of *Ly6c2* and *Itgal* (Fig. 1B and Fig. 5, A-D). The two clusters of Ly6C^neg^ macrophages likely represent resident populations of distinct hematopoietic origins (*46*). Interestingly, the Mono/Mac 4 cluster had not been previously identified in the SkM and resembles a population of infiltrating non-classical monocytes recently described in cancer models (*47*), distinguished by the expression of *Ace*, *Pou2f2*, *Adgre4* and *Treml4* (Fig. 1B and fig. S6A). In aged n.i. SkMs, we observed a remodeling of the macrophage compartment that could be mostly attributed to an expansion of the Mono/Mac 4 population (Fig. 5, A-C). To validate and interpret these findings, we quantified, by FC, the abundance of these three populations in n.i. SkM of yg and old animals. Resident macrophages were identified as CD11b^pos^Ly6C^neg^ and discriminated into two populations of Lyve1^pos^CCR2^neg^ or Lyve1^neg^CCR2^pos^. The Mono/Mac 4 population was identified in an independent panel as CD11b^pos^Ly6G^neg^Cd11a^pos^ (fig. S6, B and C). Indeed, we observed a 2,6-fold expansion of the Mono/Mac 4 population in aged n.i. SkMs (Fig. 5E), while the resident populations were either not affected (Fig. 5E, Ccr2-resident Macs) or only modestly affected (Fig. 5E, Lyve1-resident Macs, 1,4-fold increase), supporting the remodeling of the macrophage compartment in aged SkMs predicted by the scRNAseq data.

Notably, at 1 dpi, the dominant population of *Ly6c2*-expressing Mono/Macs, classically defined as a uniform population of infiltrating monocyte-derived macrophages with a pro-inflammatory profile (*14*), was composed, in fact, by three populations (Fig. 5, A-C). One population had the highest levels of *Ly6c2* and expressed genes of classical tumor and tissue-infiltrating monocyte-derived macrophages (*Vcan*, *Sell*, *Thbs1*, (*47, 48*)), along with genes expressed in peripheral monocyte-derived macrophages (*F10*, *C3*, *Sell*, *Hp*, (*49*)). This population likely corresponds to the basal state of cells of the monocytic lineage that infiltrate the SkM early after injury, and was named infiltrating monocyte-derived macrophages (infMacs, yellow, Fig. 5, A-C and F, and fig. S7A). Interestingly, we identified 2 additional clusters that shared the expression of some genes with infMacs (*Ccr2*, *Ccr5*, *Cd14*, *Fn1, Thbs1*, *F10*), but also exhibited a distinct gene expression signature of macrophages with a regulatory profile (*Arg1*, *Il10*, *Pf4*, *Vegfa*, *Spp1*, *Pdpn*, *Trem2, Hbegf*), characterized by their immunosuppressive activity in cancer models and organ transplantation (*47, 50-53*), named regeneration-associated regulatory macrophages (regMacs, blue, Fig. 5, A-C and F, and fig. S7A). Even though regMacs had a distinct immune suppressive profile, they also co-expressed high levels of common inflammatory mediators (fig. S7B) and chemoattractant factors (fig. S7C), suggesting a mixed activation state. regMacs 1 was the dominant population, while regMacs 2, less abundant, were negative for some of the markers, potentially representing a transition state (Fig. 5F and fig. S7, A-C). Importantly, the balance between infMacs and regMacs 1 was altered in aging, with an expansion of infMacs and simultaneous contraction of the regMacs 1 population (Fig. 5C).

At later stages of SkM regeneration (3 and 5 dpi), our analysis revealed the existence of four different cell states within the macrophage compartment, one of them only present in aged animals (Fig. 5, A-C). In agreement with previous reports, these populations expressed lower levels of *Ly6c2* and higher levels of *Cx3cr1* than macrophages present at 1dpi (Fig. 5F and fig. S8A). However, gene expression profiles revealed a previously unrecognized heterogeneity, challenging the idea of an unique and uniform Ly6C^low^ population of macrophages coordinating the reparative phase of SkM regeneration. One cluster, which we named repair-associated macrophages (repMacs, green, Fig. 5, A-C), was contracted from 3 to 5 dpi, and had the highest expression of genes usually associated with macrophages with pro-repair activity (Fig. 5F and fig. S8B), including growth factors (*Igf1*, *Vegfa*, *Pdgfa*), genes involved in lipid metabolism (*Fabp5*, *Fabp4*, *Lpl*) and lysosomal activity (*Ctsd*, *Ctsl*, *Ctsb*). Importantly, this cluster was selectively lost in aged SkMs at later stages of regeneration (Fig.5, B and C). A second cluster of macrophages was expanded from 3 to 5 dpi, representing a previously uncharacterized macrophage state in the SkM, that accumulates at later stages of regeneration, and was thus named late regeneration-associated macrophages (lateMacs, red, Fig. 5, A-C). This macrophage state was characterized by the expression *Tomt*, major histocompatibility complex class II (MHC II) genes (*H2-Ab1*, *H2-Aa*, *H2-Eb1,* Fig. 5F and fig. S8B) along with genes associated with interferon signaling (*Irf7*, *Ifit3*, *Ifitm3,* Fig. 5F and fig. S8B). Importantly, this population did not express markers of DC lineage (Fig. 1E and fig. S8C), supporting their macrophage identity, despite the gene profile similar to those found in DCs. The third cluster of macrophages remained constant from 3 to 5 dpi, expressing markers of repMacs and lateMacs simultaneously, along with intermediate levels of *Ly6c2*, suggesting an intermediate transition state, named intermediate macrophages (intMacs, orange, Fig. 5, A-C (Fig. 5F and fig. S8 A, B). Notably, our analysis also identified a population of macrophages that was only detected at 5 dpi in aged animals, which we named age-associated macrophages (ageMacs, black, Fig. 5, A-C). ageMacs also had a mixed expression profile, but distinctively upregulated the expression of several chemokines (*Cxcl10*, *Ccl8*, *Ccl5,* Fig. 5F and fig. S8B), which may result in a delayed resolution of the inflammatory response in aged SkMs. Moreover, since they also express a pro-fibrogenic gene signature (*Gpnmb*, *Spp1*, *Syngr1*, *Lgals3,* fig. S8D) described in dystrophic SkMs (*54*) and also present in the transient repMac population, their accumulation at 5dpi may be linked to increased fibrosis in aged SkMs.

To validate the full complexity of macrophage states during SkM regeneration identified in of our scRNAseq analysis, we developed a FC panel that would allow us to distinguish infMacs, regMacs, intMacs, repMacs and lateMacs, based on the expression profiles extracted from our scRNAseq dataset. We used Ly6C levels to discriminate infMacs and regMacs from repMacs and late Macs, along with MHC II and CD109 as lineage specific markers, (Fig. 6A and fig. S9, A and B). The identity of the macrophage populations was confirmed based on hallmark gene expression evaluated by qPCR after FACS (Fig. 5F and fig. S9 C and D). FC analysis on a time-course following SkM injury in yg animals validated the dynamics of the different macrophage populations predicted by our scRNAseq dataset (Fig. 6B and Fig. 5C). Importantly, analysis of these macrophage populations in old versus yg animals, confirmed the age-related selective loss of regMacs and repMacs during SkM regeneration (Fig. 6C).

**Fig. 6.**
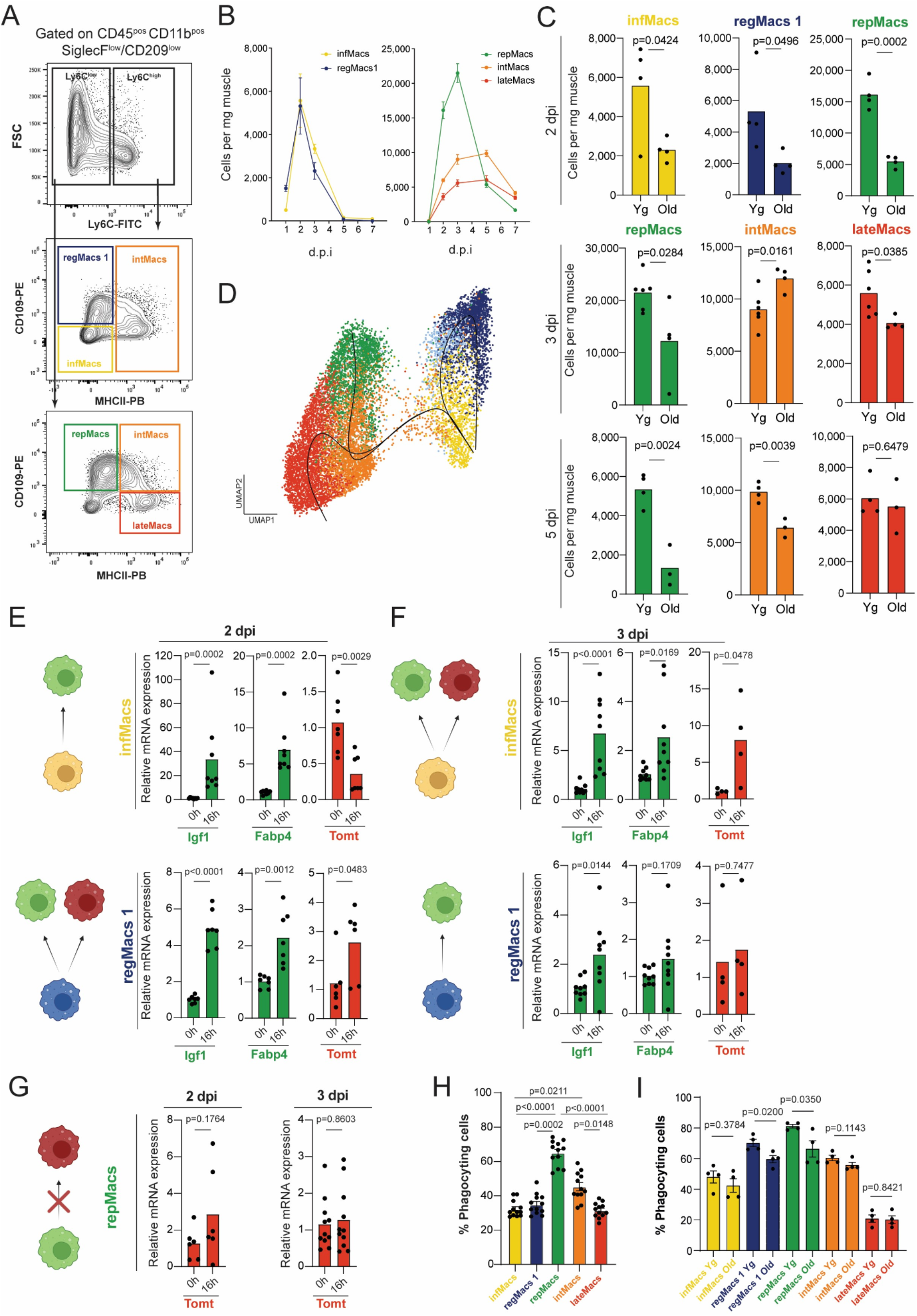
Characterization of novel macrophage populations during skeletal muscle regeneration. **A,** Gating strategy used in the flow cytometry analysis of macrophage populations during SkM regeneration gated on CD45^pos^CD11b^pos^SiglecF/CD209^low^cells: infMacs (Ly6C^high^MHCII^neg^CD109^neg^), regMacs1 (Ly6C^high^MHCII^neg^CD109^pos^), repMacs (Ly6C^low^MHCII^neg^CD109^pos^), intMacs (Ly6C^high^MHCII^pos^CD109^pos+neg^ and Ly6C^low^MHCII^pos^CD109^pos^) and lateMacs (Ly6C^low^MHCII^pos^CD109^neg^). The FMO density plots to define the positive populations are shown in fig. S9B. **B,** Quantification, by flow cytometry, of macrophage populations in regenerating SkMs of yg (2–4 months) wt (C57BL/6) mice at different timepoints following injury (n=6 for 3 dpi; n=3 for 7 dpi and n=4 for all other conditions). Gating strategy used for analysis at 3, 5 and 7 dpi is as shown in A. For the analysis at 1 dpi and 2 dpi a negative selection for Ly6G^pos^ cells was also performed to exclude neutrophils. **C,** Quantification, by flow cytometry, of macrophage populations in regenerating SkMs of yg (2–3 months) and old (24-31 months) wt (C57BL/6) mice, at 2 dpi, 3 dpi and 5 dpi (n=4 for yg and old mice at 2 dpi; n=6 for yg mice at 3dpi; n=4 for old mice at 3 dpi; n=4 for yg mice at 5 dpi and n=3 for old mice at 5 dpi); Gating strategy used for analysis at 3 and 5 dpi is as shown in A. For the analysis at 2 dpi a negative selection for Ly6G^pos^ cells was also performed to exclude neutrophils. **D,** Trajectories of macrophage populations in regenerating SkMs of yg wt mice, at 1, 3, and 5 dpi, using pseudotime values estimated from UMAP coordinates of the scRNAseq dataset with Slingshot. Cell colours in the plot follow their respective annotation (yellow - infMacs; dark blue - regMacs 1; light blue - regMacs 2; green - repMacs; orange - intMacs; red - lateMacs). We defined the trajectory starting point based on Ly6c2 expression, a marker highly expressed in the initial infiltrating populations (infMacs). **E,** Relative levels of Igf1, Fabp4 and Tomt mRNA, detected by RT-qPCR, in infMacs and regMacs1 isolated, by FACS, from 2 dpi SkMs of yg wt (C57BL/6) mice, after 0h and 16h of *in vitro* culture (n=8 for Igf1 and Fabp4 analysis in infMacs; n=6 for Tomt analysis in regMacs1; n=7 for all others analysis). **F,** Relative levels of Igf1, Fabp4 and Tomt mRNA, detected by RT-qPCR, in infMacs and regMacs isolated, by FACS, from 3 dpi SkMs of yg wt (C57BL/6) mice, after 0h and 16h of *in vitro* culture (n=9 for Igf1 and Fabp4 analysis in infMacs and regMacs1; n=4 for Tomt analysis in infMacs and regMacs1). **G,** Relative levels of Tomt mRNA, detected by RT-qPCR, in repMacs isolated, by FACS, from 2 and 3 dpi SkMs of yg wt (C57BL/6) mice, after 0h and 16h of *in vitro* culture (n=6 for analysis at 2 dpi; n=11 for analysis at 3dpi). **H,** Quantification, by flow cytometry, of the percentage of cells that ingested opsonized beads, for all macrophage populations present during SkM regeneration, assayed in single cell suspensions prepared from 3 dpi SkMs of yg (3-4 months) wt (C57BL/6) mice (n=12 for all analysis). **I,** Quantification, by flow cytometry, of the percentage of cells that ingested opsonized beads, for all macrophage populations present during SkM regeneration, assayed in single cell suspensions prepared from 2 dpi SkMs of yg (2 months) and old (24-29 months) wt (C57BL/6) mice (n=4 for all analysis). In B, data are represented as average ± s.e.m. In C and E-G data are represented as average, and each dot represents one animal. In H and I, data are represented as average ± s.e.m, and each dot represents one animal. In C, p values are from two-tailed Student’s t test. In E, for Igf1, Fabp4 and Tomt analysis in infMacs, p-values are from two-tailed Mann-Whitney test, for Igf1, Fabp4 and Tomt analysis in regMacs1 p values are from two-tailed Student’s t test. In F, for Igf1 analysis in infMacs, p value is from two-tailed Mann-Whitney test, and for all others analysis, p values are from two-tailed Student’s t test. In G, p values are from two-tailed Student’s t test for analysis at 2 dpi, and from two-tailed Mann-Whitney test for analysis at 3 dpi. In H, p-values are from Kruskal-Wallis test with Dunńs post test. In I, p value for comparison of intMacs yg versus old is from two-tailed Mann-Whitney test, and for all other analysis, p values are from two-tailed Student’s t test. SkM, skeletal muscle; yg, young; wt, wild-type; dpi, days post injury; FMO, Fluorescence Minus One Control.

### Two Ly6C^low^ macrophage states emerge through independent differentiation trajectories and represent distinct functional subsets

The complexity of the macrophage landscape uncovered by our data prompted us to interrogate the lineage relation between the multiple macrophage populations. As a first approach to address this problem, we used slingshot (*55*) to infer cell lineages and pseudotimes from our single-cell gene expression data. This analysis suggested three independent trajectories across six macrophage states (Fig 6D), challenging the prevalent view on macrophage dynamics during SkM regeneration describing a unique trajectory (*14*). infMacs and regMacs were predicted as potentially interchangeable states. One lineage predicted the emergence of repMacs from the Ly6C^high^ populations through an intermediate state of intMacs. Interestingly, an alternative lineage predicted that lateMacs also derive from Ly6C^high^ populations through an independent differentiation path. To experimentally validate the existence of two independent lineages of Ly6C^low^ macrophages, we adapted a method for *ex-vivo* lineage tracing of macrophages isolated from regenerating SkMs. In this assay, specific populations of macrophages isolated from the SkM after injury are cultured for 16h and gene expression analysis at 16h vs 0h allows monitoring macrophage state transitions as they happen *in vivo*

(*23*). First, we isolated infMacs and regMacs from SkM at 2 dpi and followed their differentiation path. In both populations, we observed a down-regulation *Ly6c2*, and of their lineage-specific markers, suggesting a transition into Ly6C^low^ states (Fig. S9E). However, infMacs only up-regulated markers of repMacs, and not of lateMacs, suggesting that at 2dpi macrophage transitions happen mostly along lineage 2 (Fig. 6E). Since lateMacs increase in number between 3 and 5dpi, we tested the ability of regMacs and infMacs isolated at 3dpi to give rise to lateMacs. Indeed, up-regulation of lateMacs markers was now detected and the up-regulation of repMac genes decreased in magnitude (Fig. 6F), suggesting that lineage 2 and 3 are separated in time. Importantly, repMacs isolated at 2dpi or 3dpi did not up-regulate markers of lateMacs (Fig. 6G), confirming the independence of the two lineages.

Macrophage function during SkM regeneration has been mostly linked with clearance of necrotic debris by phagocytosis and regulation of MuSC function (*14*), both roles being affected by aging (*23-25*). Thus, we tested the phagocytic capacity of our newly identified macrophage populations. To quantify phagocytic activity, we developed an assay where single cell suspensions prepared from regenerating SkM at 3dpi were incubated with opsonized beads for 3h *ex vivo*, and bead uptake by each macrophage population was determined by FC (Fig. S10A). Surprisingly, we found that the Ly6c^low^population of repMacs displayed the highest phagocytic activity (Fig 6H, 65% of cells ingested beads), challenging the prevalent idea that Ly6C^high^ macrophages are the most phagocytic (Fig. 6H, only 35% of regMacs and 32% of infMacs ingested beads). Interestingly, lateMacs displayed similar phagocytic activity to Ly6C^high^ populations (Fig. 6H, 32% of cells ingested beads), suggesting that phagocytosis is differentially regulated along the two differentiation lineages of Ly6C^low^ macrophages, with the lineage of repMacs upregulating their phagocytic activity while transitioning into the Ly6C^low^ state. Importantly, in addition to their numerical loss, repMacs and regMacs also showed a decline in their phagocytic capacity with age (Fig. 6I).

Collectively, these data uncovered a complex landscape of macrophage activation through multiple stages of SkM regeneration, including: (i) a previously unidentified population of regulatory macrophages detected in the early stages of regeneration that is decreased in aged animals and (ii) two independent sates Ly6C^low^ macrophages, associated with the reparative phase of SkM regeneration, that differentiate along independent trajectories, are functionally distinct and differentially affected by aging.

## DISCUSSION

Age-related alterations in the tissue’s immune environment are emerging as important contributors to impairments found in aged organs (*56, 57*). However, their contribution to regenerative decline in aging is just starting to be explored (*23-25*). Here, we performed a complete analysis of the immune environment in the aged regenerating SkM, with time and single cell resolution. We uncovered previously undescribed immune cell types participating in regeneration and age-related alterations in abundance and cell states within the lymphoid and myeloid compartments, with potential to impact SkM health and regenerative capacity in aging.

The importance of immune aging for SkM homeostatic health has been demonstrated in studies using heterochronic bone marrow (BM) transplants. BM from old donors transplanted into yg hosts caused reduced MuSC numbers and promoted fibrogenic conversion (*58*). Consistently, it has been reported that macrophage ablation prevents age-related decline in SkM strength (*59*) and that BM transplants from yg mice into old donors is sufficient to ameliorate signs of sarcopenia (*58*). Despite this knowledge, the identity of the immune populations and signals mediating these effects remains mostly elusive. In our analysis, the most prominent feature of the aged immune compartment in homeostatic SkM was an expansion of lymphocytes, due to the emergence of age-specific populations of B and T cells. This phenomenon has been described in multiple aged organs, suggestive of systemic changes affecting B and T cell aging (*29*). Although the age-related factors driving the expansion of aged lymphoid subsets are still unclear, the priming of B cells has been associated with the stimulation of TLR7 and TLR9 signaling, likely caused by the increase presence of damage associated molecular patterns (DAMPs) (*33*), and maintained by IFNγ (*60*). Interestingly, in neurogenic niches of the aged brain, T cells are clonally expanded and distinct from circulatory populations, suggesting local effects due to exposure to niche-specific antigens. Similarly to our findings in the SkM, T cells from old brain express higher levels of IFNγ, causally linked to a decrease in neural stem cell proliferation (*61*) and potentially associated with the expansion of aged B cells (*60*). Moreover, mouse models of accelerated T cell aging showed signs of premature sarcopenia, including reduced fiber size and strength, and insulin resistance (*62*). In addition to lymphocytes, our analysis revealed an expansion of neutrophils, specific DC subtypes and a population of infiltrating non-classical monocytes. Also here, damage accumulated in the SkM throughout life could be the origin of the DAMPs driving the infiltration or local expansion of these immune cell types. Further work is necessary to understand the consequences of these immune alterations for the decline of SkM health in aging.

Contrasting with the generalized expansion of most immune cell types in the homeostatic SkM, the injury-associated immune response in aging was broadly characterized by defects in the accumulation of different populations, supporting and expanding previous studies (*23-25*). Interestingly, these defects are a common feature of the aged immune response after injury, as recently reported in the spinal cord (*63*). The causes of these alterations are unknown, but could potentially be associated with impairments at multiple levels, including hematopoietic response to tissue injury (*64*), or signaling axis involved in immune cell recruitment or retention at the injury site.

Eosinophils and neutrophils were among the most affected cell types in terms of cell abundance, in the early stages after injury. Given their significant contribution to SkM regeneration in yg animals (*65, 66*), these defects may be part of the mechanism through which immune aging drives regenerative decline. Similarly, injury-associated DCs, particularly, pDCs were lost from SkM of aged mice throughout the whole regeneration process. However, the role of DCs during SkM regeneration has never been studied and this is the first report of their diversity and dynamic regulation after SkM injury. Thus, to understand the consequences of age-related DC loss for regenerative decline it will be required to first determine which are the mechanisms linking DC function and regenerative efficiency, which may include direct effects on MuSCs (*67*) or classical immune regulatory functions (*68*). Interestingly, total T cell abundance and dynamics was not altered in aged regenerating SkMs. Since Tregs fail to accumulate in aged SkMs following injury (*69*), it is possible that other T cell subsets are expanded, resulting in a remodeling of the T cell compartment, rather than a numerical loss of T cells. Due to the high influx of myeloid cells following injury, our scRNAseq analysis did not capture sufficient lymphocytes from regenerating SkMs that would allow addressing this question. In the future, it would be interesting to perform a scRNAseq analysis of lymphoid enriched immune cells (CD45^pos^CD11b^neg^) in the same experimental conditions to determine whether the age-specific subsets of T cells identified in homeostasis are also present and expanded during regeneration and explore the potential contribution of T cell aging for regenerative failure.

Defects in cell accumulation in the early phase of SkM regeneration also affected the dominant populations of recruited macrophages. Surprisingly, our analysis found that this is an heterogenous population comprising two distinct macrophage states. regMacs had been identified in cancer models and in the context of organ transplantation (*47, 51*), but never in the regenerating SkM. These cells were shown to have immune suppressive activity, favoring Treg expansion and effector functions while promoting the depletion of activated T cells (*51, 70*). Thus, the loss of regMacs in aged regenerating SkMs could be a mechanism driving the proposed remodeling of the T cell compartment.

During the late phase of regeneration, we uncovered an unanticipated heterogeneity within the Ly6C^low^ macrophage population. Interestingly, a recent study analyzing CD45^pos^ cells isolated from CTX-injured SkM at 4dpi has also identified 4 macrophage subsets that share some of the characteristics reported here (*71*). The current view on the process of macrophage phenotypic transitions during SkM regeneration postulates that inflammatory macrophages give rise to a pro-repair state, characterized by the downregulation of Ly6C and the upregulation of Cx3cr1 (*14*). Instead, our analysis revealed two populations that are functionally distinct and emerge through independent differentiation trajectories. The population of repMacs resembles what is commonly associated with a pro-repair function, expressing genes involved in angiogenesis and growth factors, along with a signature of lipid metabolism and lysosomal activity, known to be involved in the process of macrophage phenotypic transition (*14*). Our functional assays showed that this population had the highest phagocytic activity, and was the most affected by aging. These results refine our previous findings showing that a defect in the accumulation in pro-repair macrophages is associated with SkM regenerative failure in aging (*23*). Furthermore, these results suggest that the accumulation of uncleared necrotic fibers in regenerating aged SkMs might be linked with the decline of this population, which is the most phagocytic, a function that is also impaired in aging.

Unexpectedly, our analyses predicted and validated an alternative differentiation trajectory leading to the emergence of a distinct Ly6C^low^ macrophage subtype. lateMacs are characterized by the expression of genes associated with antigen presentation and interferon response. Recently, one study reported the presence of interferon-responsive macrophage population affecting MuSC expansion through the secretion of CXCL10 (*72*). Interestingly, this population is also Ly6C^high^, potentially pointing to an overlap with our intMacs, a mixed population that shares the interferon-response signature with lateMacs and has the highest expression of *Cxcl10*. Although lateMacs and intMacs were only modestly affected by aging, we identified a new population of macrophages that was specific for the aged SkM in the late phase of regeneration. ageMacs share a mixed gene expression profile with intMacs, potentially corresponding to a state of arrested transition with pathogenic features. Indeed, ageMacs upregulate chemoattractant factors and maintain the fibrogenic gene signature, which may contribute to increased fibrosis and delayed resolution of inflammation after injury.

Our study expands and clarifies established notions on immune regulation of SkM regeneration, revealing an unanticipated complexity in immune populations that have been regarded as homogeneous and a profound remodeling of the immune compartment associated with regeneration during aging. These discoveries strengthen the idea that immune modulation could be an attractive strategy to improve regenerative capacity in aging, and provide a new set of targets to develop these immune modulatory interventions.

## MATERIALS AND METHODS

### Animals

All mice used in these studies were housed at the Direção Geral da Alimentação e Veterinaria (DGAV) accredited rodent facility of Instituto de Medicina Molecular, in individually ventilated cages within a specific and opportunistic pathogen-free (SPOF) facility, on a standard 12/12 h light cycle. All mouse work reported in this study complied with relevant institutional and national animal welfare laws, guidelines and policies. The experimental protocol was reviewed and approved by iMM ORBEA and licensed by DGAV (project license number 022860\2020). Old wild-type C57BL/6 mice were purchased from Charles River, Europe, at the age of 18–20 months and further aged in-house until analysis. Yg wild-type mice were either purchased from Charles River, Europe, or born in-house and generated using C57BL/6 breeders purchased from Charles River, Europe. Female and Males animals were used in the studies reported.

### Induction of SkM regeneration

Regeneration of SkM was induced by intramuscular injection of 50 μl of sterile 1.2% barium chloride (Sigma: 342920) in sterile saline solution (0.9% NaCl, B. Braun) into the quadriceps muscles of the mice. Mice were euthanized, at the designated time points after injury, and muscles were collected for analysis. Animals were anesthetized for the procedure with isoflurane inhalation.

### Fluorescence-activated cell sorting (FACS) and flow cytometry (FC) analysis

Single cell suspensions were obtained from dissected muscles mechanically disrupted and enzymatically digested at 37 °C for 1h using DMEM 1% P/S medium containing 0.2% collagenase B (Roche) and Calcium dichloride (CaCl_2_) 0.5 mM, and then filtered through 70 µm cell strainers (Falcon). Cells were incubated for 10 min on ice in 1x Red Blood Cell (RBC) lysis buffer and resuspended in 1ml of DMEM 10% FBS 1% P/S medium for counting in a Neubauer Chamber. Single cell suspensions were then incubated with fluorochrome-conjugated antibodies in PBS 5% HS at a density of 1×10^6^ cells/100μl, for 30 minutes at 4°C, protected from the light. Information about the antibodies used is presented in table S1. To identify viable cells, LIVE/DEAD Fixable Near-IR Dead Cell Stain (Invitrogen) was also included at a 1:1000 dilution.

The isolation of cell populations by FACS was performed at the Flow Cytometry facility of Instituto de Medicina Molecular using a FACSAria III (BD Bioscience) or FACSAria Fusion (BD Bioscience) with the software FACSDiva 6.1.3. For single cell RNA-seq analysis, CD45^pos^viable cells were sorted. For further RT-qPCR validation studies, viable CD45^pos^CD209^High^ DCs were sorted and distinct populations of macrophages were isolated based on the gating strategy presented in Fig. 6A.

FC analyses of SkM single cell suspensions were performed with the cell analyser LSRFortessa X-20 (BD Bioscience) using the FACSDiva 8.0 software. The FC data were analysed on FlowJo (BD Biosciences) software. Information about the gating strategies used for the characterization of lymphocytes, DCs, granulocytes and macrophages subtypes are present in the supplementary figs. S3, S4, S5 and Fig. 6, respectively.

### Single-cell RNA sequencing by 10× Genomics platform

#### Single cell library preparation

CD45^pos^ cells were isolated by FACS, from n.i. and injured SkM at 1, 3 and 5 dpi from yg (3 months) and old (25 months) female mice. Cells were isolated in two sessions using yg and old mice: n.i. and injured SkMs at 3 dpi in the first session, injured SkMs at 1 dpi and 5 dpi in the second session. For each condition, isolated CD45^pos^ cells from two independent animals were pooled as one sample. After FACS, cell concentration and viability (80-90%, see fig. S1B) of each cell suspension were determined using 0.4% trypan blue (Gibco™: 15250061). Cells were washed and resuspended in 1x PBS (calcium and magnesium free) containing 0.04% BSA, to a final concentration of 1000 cells/µl. In brief, 20,000 CD45^pos^ cells per condition were loaded onto a 10X Genomics Chromium platform for Gel Bead-in-Emulsion (GEM), complementary DNA (cDNA) generation was performed, and sequencing libraries created with cell- and transcript-specific barcodes, using the Chromium Next GEM Single Cell 3’ Kit v3.1 (10× Genomics: PN-1000268), Chromium Next GEM Chip G Single Cell Kit (10× Genomics: PN-1000127) and the Dual Index Kit TT Set A (10× Genomics: PN-1000215), following the supplier’s protocol. The quality of cDNAs were assessed (see fig. S1B) on LabChip® GX Touch™ Nucleic Acid Analyzer, using DNA High Sensitivity Reagent Kit (PerkinElmer: CLS760672) and LabChip GX Touch 24 (PerkinElmer: CLS138948). Libraries were sequenced using the Illumina NovaSeq6000 PE150 targeting 20,000 reads per cell, using services at Novogene (Cambridge, UK).

#### Sequencing data analysis

##### Read mapping, quality control and library size normalization

The feature-barcode matrices were obtained by aligning reads to the mouse genome (GENCODE vM23/Ensembl 98) using CellRanger v7.0.1. We initially identified a total of 65,006 cells. All sample data were imported into R (version 4.1.2), merged and processed together. We performed quality filtering by retaining only cells that met the following criteria: a minimum library size of 3162 UMIs, a minimum of 3000 expressed genes and mitochondria gene content below 10% (Fig. S1 C-E). We identified doublets using the scDblFinder package (version 1.8.0, https://bioconductor.org/packages/release/bioc/html/scDblFinder.html) and excluded them from the dataset. The final dataset consisted of 40,956 cells. We used the scran package (version 1.22.1, https://bioconductor.org/packages/release/bioc/html/scran.html) to perform normalization of library size biases due to differences in sequencing depth per cell.

##### Dimensionality reduction and clustering analysis

We performed the following analysis using the Seurat package (version 4.1.1, https://satijalab.org/seurat/) unless mentioned otherwise. Firstly, we selected the top 3000 highly variable genes (HVGs) which we identified using the FindVariableFeatures with the selection.method “vst”. We conducted principal component analysis (PCA) and Uniform Manifold Approximation and Projection for Dimension Reduction (UMAP) using the HVGs. Subsequently, we utilized the top 25 principal components (PCs) for constructing t-SNE plots. We performed shared nearest neighbor (SNN) modularity optimization-based clustering using the top 30 PCs. In order to identify biologically meaningful clusters, we used a window of resolutions from 0.1 to 1. Using the package clustree (version 0.5.0, https://cran.r-project.org/web/packages/clustree/index.html) we constructed clustering trees showing the relationship between the clusters at different resolutions (Fig. S11). This allows us to inspect which clusters are clearly distinct and which are unstable. Final assessment was performed by evaluating the expression of specific markers in each cluster.

##### Differential expression analysis

We performed differential gene expression analysis on the normalized dataset using the FindAllMarkers/FindMarkers function from the Seurat package. We tested for differential gene expression of the clusters (cluster-based analysis) on each subset using the MAST differential expression test. We restricted testing to genes that exhibited on average a minimum 0.25-fold difference (log-scale) between the two groups. We excluded clusters with less than 40 cells and compared the cluster of interest against all the remaining clusters.

##### Macrophage subset

For each macrophage subset including either all populations or only yg samples, we performed a reiteration of the upstream analysis pipeline. Furthermore, batch effect correction was performed using the limma package (version 3.50.3, https://bioconductor.org/packages/release/bioc/html/limma.html). We used the removeBatchEffect function for removing unwanted batch effects, associated with the technical variables of “sequencing lane” and “sorting day”.

##### Gene Set Enrichment Analysis

We used the fgsea package (https://github.com/ctlab/fgsea) for the pathway enrichment analysis of T cells in homeostatic muscle prior to injury. We performed the MAST test for differential gene expression implemented in Seurat on the homeostatic muscle T cells. We set the thresholds for log-fold change (logfc.threshold) and fraction of cells expressing the gene (min.pct) to 0 to minimize genes excluded from the analysis and ranked the genes on their log-fold change values. We then provided this ranked list of genes alongside the hallmark pathways (Mouse MSigDB Collections https://www.gsea-msigdb.org/gsea/msigdb/mouse/collections.jsp) as inputs to the ‘fgsea’ function in R to compute the normalized enrichment score (NES) for each hallmark pathway. We excluded pathways with gene set size below 10 and above 500 using the arguments ‘minSize’ and ‘maxSize’ to set the thresholds. We then filtered the results for pathways with NES higher than 1 or lower than -1.

##### Cell-cell communication analysis

We used the create CellChat function from the CellChat R package (*36*) to assess cell-cell communication by identifying ligand-receptor (LR) pairs with altered expression in Old vs Yg samples. We performed a separate analysis for each combination of time point and aging condition, using the clusters that were considered for the upstream differential gene expression analyses (with over 40 cells).

We adapted the workflow provided by the CellChat development group ((*36*), https://github.com/sqjin/CellChat/). We first identified overexpressed genes that code for LR pairs between cell types based on the results of the cluster-based differential gene expression analysis using the function identifyOverExpressedInteractions. We then computed communication probabilities using the computeCommunProb function. We used the netVisual_heatmap function to observe the differential number of interactions and interaction strength of B cells as signal senders in homeostatic SkM prior to injury (Fig. S3G). To identify individual LR pairs in each cell group of interest, we used the netMappingDEG function in order to map the genes differentially expressed with age across the LR pairs in the CellChat database as either a ligand or a receptor of the pair. We then separated interactions based on the interaction type annotation provided by the database and on whether the genes involved in the interaction are up or downregulated, and used the netVisual_chord_gene function in order to visualize the chord diagram of the identified LR pairs, with B cells as the ‘sources.use’ argument and T, B, and NK cells as the ‘targets.use’ argument (Fig. 2H).

##### Trajectory inference analysis

For trajectory inference analysis, we used the macrophage population subset including only yg samples. We then computed trajectories across the clusters using the Slingshot package ((*55*), https://www.bioconductor.org/packages/release/bioc/html/slingshot.html) using the 10 UMAP components constructed using the normalized log-scale count matrix (without batch effect correction). We defined a starting point for the trajectory curves based on the highest expression of Ly6c2, a marker that has highest expression in the initial infiltrating populations, then computed the pseudotime values using the slingshot function, and constructed the curve for each trajectory to display on UMAP coordinates using the SlingshotDataSet function.

### Ex vivo macrophage analysis

Subtypes of macrophages from 2 dpi and 3 dpi SkMs, were isolated by FACS as described above, and 20.000-50.000 cells were collected for analysis at 0 h or cultured and collected for analysis after 16 h. Cells were incubated at 37 °C in DMEM supplemented with 10% FBS and 1% P/S. Collected cells were used for RNA analysis.

### Phagocytosis assays in whole SkM single cell suspensions

Evaluation of phagocytic uptake was performed using Fluoresbrite® BB Carboxylate Microspheres 1.75µm (Polysciences, 17686-5). Opsonization of the microspheres was performed by incubation in 50% FBS in PBS for 30 min at 37°C. Opsonized microspheres were added to 500.000 cells from whole SkMs single cell suspensions, obtained as described above, at a concentration of 3.4 × 10^8^ particles per ml, the mixture was plated in 6-well plates in DMEM supplemented with 10% FBS and 1% P/S and incubated for 3 h at 37°C.

Phagocytic uptake was measured by FC through the quantification of the percentage of each macrophage population that was positive for BB (Bright Blue) fluorophore. Macrophage populations were defined using the gating strategy in Fig. 6A, adapting the fluorophores used to allow compatibility with the BB fluorophore conjugated with the microspheres. Gating strategy to detect BB fluorescence in each macrophage population and corresponding FMOs to define the phagocytic fraction is shown in fig. S10.

### RNA isolation and quantitative real-time PCR assay

Total RNA from cells isolated by FACS was extracted using an RNeasy Micro kit (Qiagen) according to the supplier’s instructions.

cDNA was synthesized using an iScript cDNA synthesis kit (BioRad). Real-time PCR was carried out on a ViiA 7 Real-Time PCR System (Thermofisher Scientific), using PowerUp SYBR Green Master Mix (Applied Biosystems). Specific gene expression in each sample was normalized to β-actin levels, and results are presented as gene expression levels relative to levels in control samples, which were arbitrarily set to one. For information on primer sequences, see table S2.

### Statistic and reproducibility

For the experiments reported, a minimum of three independently manipulated animals were used per condition. In the graphs reporting data collected from animals, each dot corresponds to an independent measurement from an independent animal. For figures reporting *ex vivo* assays, each dot in the graph corresponds to an independent measurement from an independent assay in a culture generated from an independent animal or pool of two animals (when cell numbers were limiting). No statistical methods were used to predetermine sample sizes, but our sample sizes were similar to those reported in previous publications (*23, 24*). No randomization method was used to allocate animals to experimental groups. Data collection and analysis were not performed blind to the conditions of the experiments.

For data from FC and RT-qPCR analysis, results are presented as average and s.e.m. and statistical analysis was carried out using GraphPad Prism 9 and all data sets were tested for a normal distribution with Shapiro-Wilk test before proceeding with parametric or non-parametric analyses. For comparisons between two groups, statistical significance was tested using two-tailed Student’s t test for data sets with normal distributions or with the two-tailed Mann–Whitney test for data sets without a normal distribution. For multiple comparisons, statistical significance was tested using one-way ANOVA with Tukey post test for data sets with normal distributions or with the Kruskal-Wallis test with Dunńs post test for data sets without a normal distribution.

Statistical methods used for the single-cell RNA sequencing analysis are described in detail in the sections above.

## ACKNOWLEDGMENTS

We would like to thank the Flow Cytometry and Rodent facilities of Instituto de Medicina Molecular João Lobo Antunes for their technical support, and Ana Espada Sousa and Luis Graça for the critical reading and suggestions on the manuscript.

## Funding

European Molecular Biology Organization Installation Grant IG4448 (PSV)

Fundação para a Ciência e Tecnologia grant PTDC/MED-OUT/8010/2020 and EXPL/MED-OUT/1601/2021 (PSV and JN).

Fundação para a Ciência e Tecnologia contract CEECIND/00436/2018 (NLBM) and 2021.03843.CEECIND (JN).

Fundação para a Ciência e Tecnologia fellowships 2022.14294.BD (NSS), 2020.05627.BD (MB), UI/BD/154081/2022 (TC) and 2022.10705.BD (IBM)

## Author contributions

Conceptualization: PSV, JN

Experimental design: NSS, PSV, JN

Investigation: NSS, MB, MFB, IBA, IAE, TC, IBM

Data analysis: NSS, MB, MFB, TC

Supervision: NLBM, PSV, JN

Writing—original draft: PSV, JN

Writing—review & editing: NSS, MB, TC, NLBM

## Data and materials availability

All the data generated or analysed during this study are included in the published article and its Supplementary Information, and are available from the corresponding author upon reasonable request. scRNAseq data generated will be available on the NCBI Gene Expression Omnibus database by the publication date

Correspondence and requests for materials should be addressed to PSV and JN.

