## Supplementary Materials for "The immune landscape of murine skeletal muscle regeneration and aging"

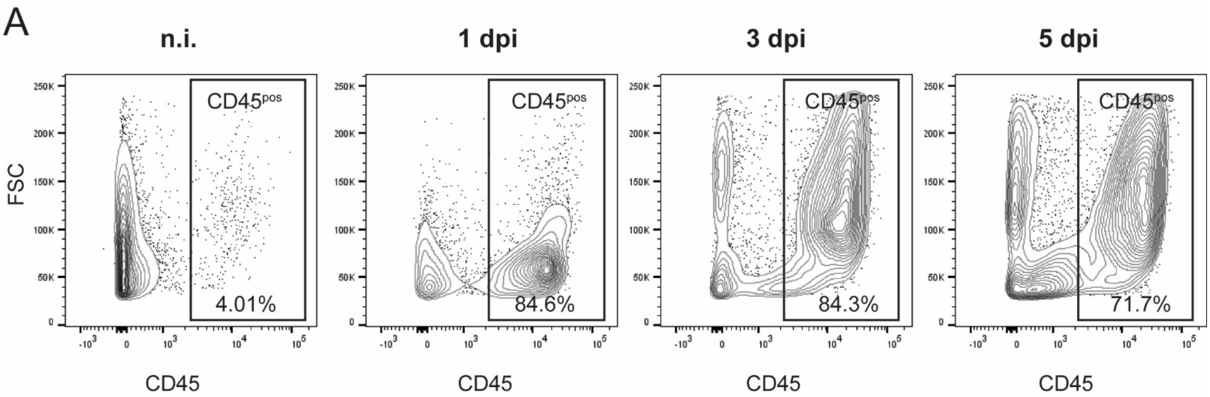


B

| Sample ID | Condition | Age | % cell viability | 1st Quality control | 2nd Quality control | | | |
|---|---|---|---|---|---|---|---|---|
| | | | | Pre library cDNA concentration (200-5000 bps) (ng/ul) | Post library cDNA concentration (ng/ul) | Average fragment size (bp) | i7 Index Sequences | i5 Index Sequences |
| **YG0** | non injured | 2 mo | 87 | 0.79 | 6.92 | 640 | GTAACATGCG | AGTGTTACCT |
| **OLD0** | non injured | 25 mo | 88 | 0.84 | 9.12 | 652 | GTGGATCAAA | GCCAACCCTG |
| **YG3** | 3dpi | 2 mo | 88 | 1.02 | 8.39 | 628 | CACTACGAAA | TTAGACTGAT |
| **OLD3** | 3dpi | 25 mo | 86 | 0.85 | 13.53 | 811 | CTCTAGCGAG | TATCTTCATC |
| **YG1** | 1dpi | 2 mo | 91 | 0.73 | 17.78 | 415 | GTAGCCCTGT | GAGCATCTAT |
| **OLD1** | 1dpi | 25 mo | 78 | 1.07 | 16.39 | 459 | TAACGCGTGA | CCCTAACTTC |
| **YG5** | 5dpi | 2 mo | 90 | 1.09 | 9.45 | 532 | TCCCAAGGGT | TACTACCTTT |
| **OLD5** | 5dpi | 25 mo | 80 | 0.66 | 13.7 | 477 | CGAAGTATAC | GAACTTGGAG |

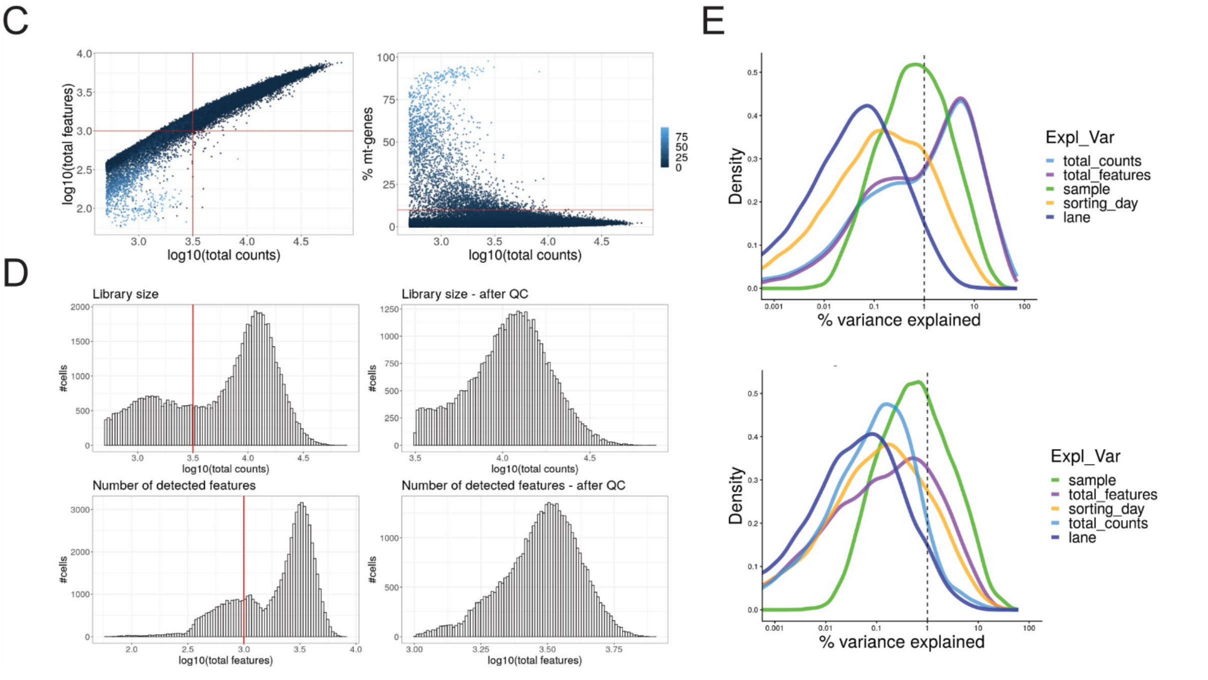


**Fig. S1| Sequencing metrics and quality control measures for the libraries obtained from samples of Cd45$^{pos}$ cells from skeletal muscle during regeneration.**

**A,** Representative density plots showing CD45$^{pos}$ cells gated on single cells, in n.i, 1, 3 and 5 dpi SkMs of yg wt (C57BL/6) mice. Percentage of CD45$^{pos}$ cells in each condition is shown.

**B,** Summary of sample information and quality control measures for the single cell library preparation.

**C,** Scatterplots showing the quality control metrics used to filter poor quality cells: total number of UMIs (total counts), number of unique features detected (total features) and the percentage of reads aligned to mitochondrial genes in each cell (% mt-genes ). Plots are colored by the % mt-genes. The filter cut-offs are indicated by the red lines.
**D,** Histograms of the library sizes (total counts, top panels) and the number of unique features detected (total features, bottom panels) in all cells of the dataset. The filter cut-offs are indicated by the red lines.
**E,** Density plots showing the percentage of the variance of expression values that is explained by the set of known variables, before (top) and after (bottom) normalization of library sizes.
SkM, skeletal muscle; yg, young; wt, wild-type; n.i., non-injured; dpi, days post injury; FSC, Forward Scatter; mo, months.

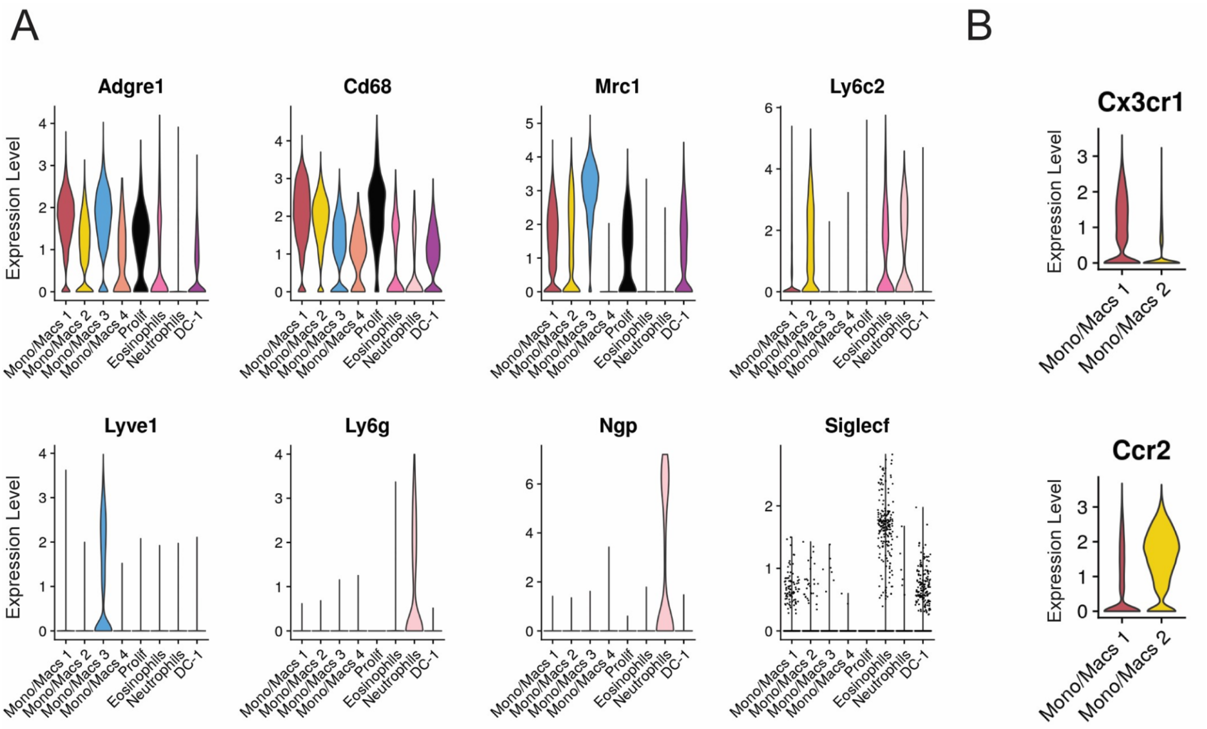


**Fig. S2| Distributions of marker gene expression levels across distinct clusters of myeloid cell types.**
**A,** Violin plots showing normalized and log-transformed expression of representative markers across clusters of myeloid cell types represented in Fig. 1D.
**B,** Violin plots showing normalized and log-transformed expression of Cx3cr1 and Ccr2 across clusters Mono/Macs1 and 2 represented in Fig. 1D.
Mono, monocyte; Macs, macrophages; DCs, dendritic cells; Prolif, proliferating cells.

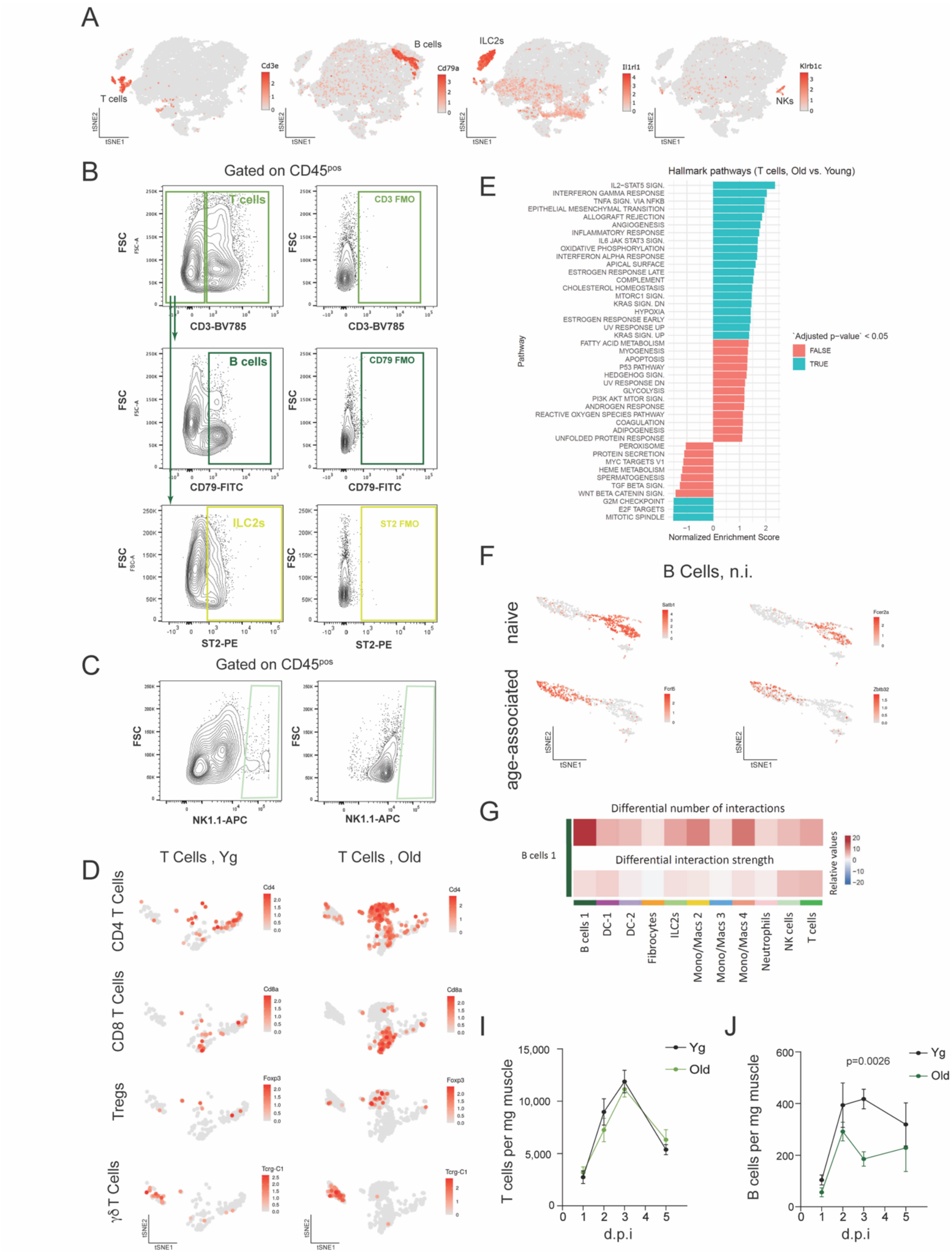


**Fig. S3 | The lymphoid compartment in homeostatic skeletal muscle of yg and old mice**
**A,** tSNE plot showing the normalized and log-transformed expression levels of Cd3e, CD79a, Il1rl1, Klrb1c genes in the whole immune compartment captured by the scRNAseq analysis. Compare with Fig. 1D to match markers with their corresponding lymphoid subtypes.

**B,** Gating strategy used in the flow cytometry analysis of lymphocytes in SkM, gated on CD45$^{pos}$ cells: T cells (CD3$^{pos}$), B cells (CD3e$^{neg}$CD79$^{pos}$), and ILC2, (CD3$^{neg}$ST2$^{pos}$). The FMO density plots displayed were used to define the positive populations.
**C,** Gating strategy used in the flow cytometry analysis of NKs in SkM, gated on CD45$^{pos}$ cells: NKs (Nk1.1$^{pos}$), gated on CD45$^{pos}$ cells. The FMO density plot displayed was used to define the positive population.
**D,** tSNE plot showing the normalized and log-transformed expression levels of Cd4, Cd8a, Foxp3 and Tcrg-C1genes in the T cell compartment.
**E,** Normalized enrichment score (NES) of MSigDB hallmark pathways between yg (control) and old T cells in n.i. SkMs of wt mice.
**F,** tSNE plot showing the normalized and log-transformed expression levels of Fcer2a, Satb1, Fcrl5 and Zbtb32 in the B cell compartment.
**G,** CellChat heatmap showing variations in number and strength of the interactions exerted by B cells (vertical axis) over other immune cell types (horizontal axis) in old vs yg (control) n.i. SkMs. Positive values (red) represent an overall increase in the interaction number/strength between B cells and a given cell type, while negative values (blue) represent an overall decrease in the interaction number/strength.
**H,** Quantification, by flow cytometry, of T cells, in regenerating SkMs of yg (3–4 months) and old (27–30 months) wt (C57BL/6) mice at different time points following injury (n=7 for yg mice at 1 dpi; n=8 for yg mice at 2 dpi; n=5 for yg mice at 5 dpi; n=3 for old mice at 1dpi; n=4 for all other conditions).
**I,** Quantification, by flow cytometry, of B cells, in regenerating SkMs of yg (3–4 months) and old (27–30 months) wt (C57BL/6) mice at different time points following injury (n=7 for yg mice at 1 dpi; n=8 for yg mice at 2 dpi; n=5 for yg mice at 5 dpi; n=3 for old mice at 1dpi; n=4 for all other conditions). The p value displayed refers to the 3dpi data and is from two-tailed t-Student test.
SkM, skeletal muscle; yg, young; wt, wild-type; n.i., non-injured; dpi, days post injury; NKs, Natural Killer cells; ILC2s, Innate Lymphoid cells type 2; FSC, Forward Scatter; FMO, Fluorescence Minus One Control.

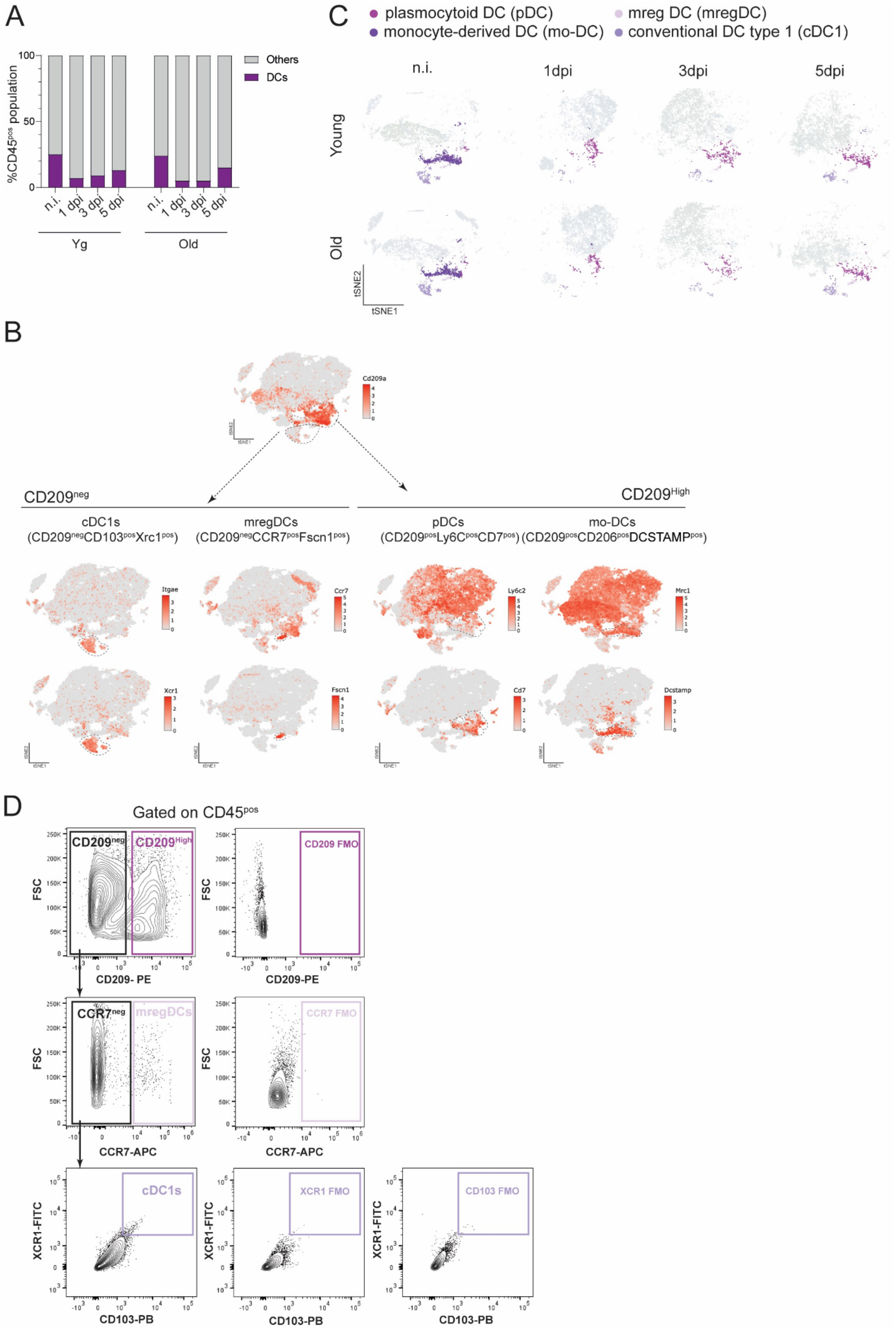

A
B
C
D
Others
DCs
%CD45pos population
n.i.
1 dpi
3 dpi
5 dpi
Yg
Old
plasmocytoid DC (pDC)
monocyte-derived DC (mo-DC)
mreg DC (mregDC)
conventional DC type 1 (cDC1)
1dpi
3dpi
5dpi
Young
tSNE2
tSNE1
Cd209a
CD209neg
CD209High
cDC1s
(CD209negCD103posXrc1pos)
mregDCs
(CD209negCCR7posFscn1pos)
pDCs
(CD209posLy6CposCD7pos)
mo-DCs
(CD209posCD206posDCSTAMPpos)
Itgae
Ccr7
Ly6c2
Mrc1
Xcr1
Fscn1
Cd7
Dcstamp
Gated on CD45pos
CD209 FMO
FSC
CD209- PE
CCR7neg
mregDCs
CCR7 FMO
CCR7-APC
cDC1s
XCR1 FMO
CD103 FMO
XCR1-FITC
CD103-PB

**Fig. S4| Discrimination of dendritic cell subtypes in skeletal muscle at homeostasis and during regeneration.**

**A,** Relative proportions of the total DC population in the $CD45^{pos}$ immune compartment identified by the scRNAseq analysis, in n.i. and injured (1, 3 and 5 dpi) SkMs of yg and old mice.

**B,** tSNE plot showing the normalized and log-transformed expression levels of genes of interest to discriminate four unique DCs subtypes in the whole immune compartment captured by the scRNAseq analysis.

**C,** tSNE plots (0.4 resolution) displaying the four unique clusters of DC populations identified in SkMs as shown in Fig. 3A, separated by time point and age. Colors represent clusters of subtypes of DCs. Non-DC clusters are shown in grey.

**D,** Gating strategy used in the flow cytometry analysis of DCs in SkM, gated on $CD45^{pos}$ cells: moDCs, in n.i. SkM and pDCs in injured SkM ($CD209^{high)}$, mregDCs ($CD209^{neg}CCR7^{pos}$), and cDC1s ($CD209^{neg}CCR7^{neg}XCR1^{pos}$ $CD103^{pos}$). The FMO density plots displayed were used to define the positive populations.

SkM, skeletal muscle; yg, young; wt, wild-type; n.i., non-injured; dpi, days post injury; DCs, dendritic cells; moDCs, monocyte-derived dendritic cells; pDCs, plasmacytoid dendritic cells; cDC1s, conventional dendritic cells type 1; mregDCs, mature dendritic cells enriched in immunoregulatory molecules; FSC, Forward Scatter; FMO, Fluorescence Minus One Control.

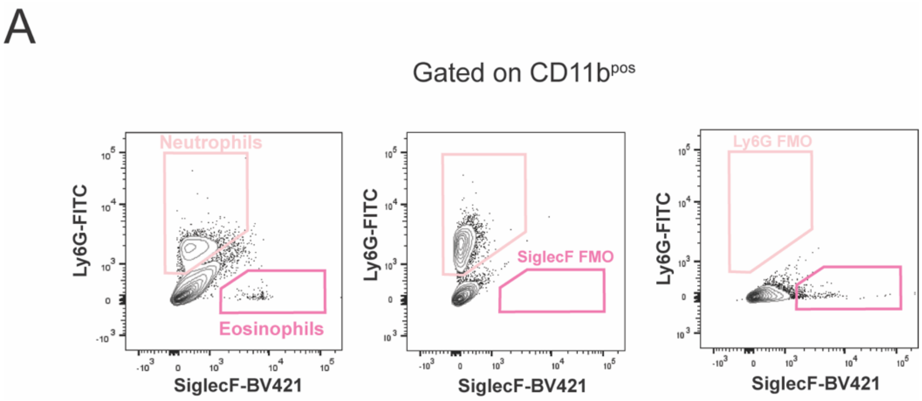


**Fig. S5 | Flow cytometry analysis of granulocytes in homeostatic and regenerating skeletal muscle**

**A,** Gating strategy used in the flow cytometry analysis of neutrophils and eosinopfils in SkM, gated on $CD11b^{pos}$ cells: Neutrophils ($Ly6G^{pos}SiglecF^{neg}$) and eosinophils ($Ly6G^{neg}SiglecF^{pos}$) The FMO density plots displayed were used to define the positive populations.

FMO, Fluorescence Minus One Control.

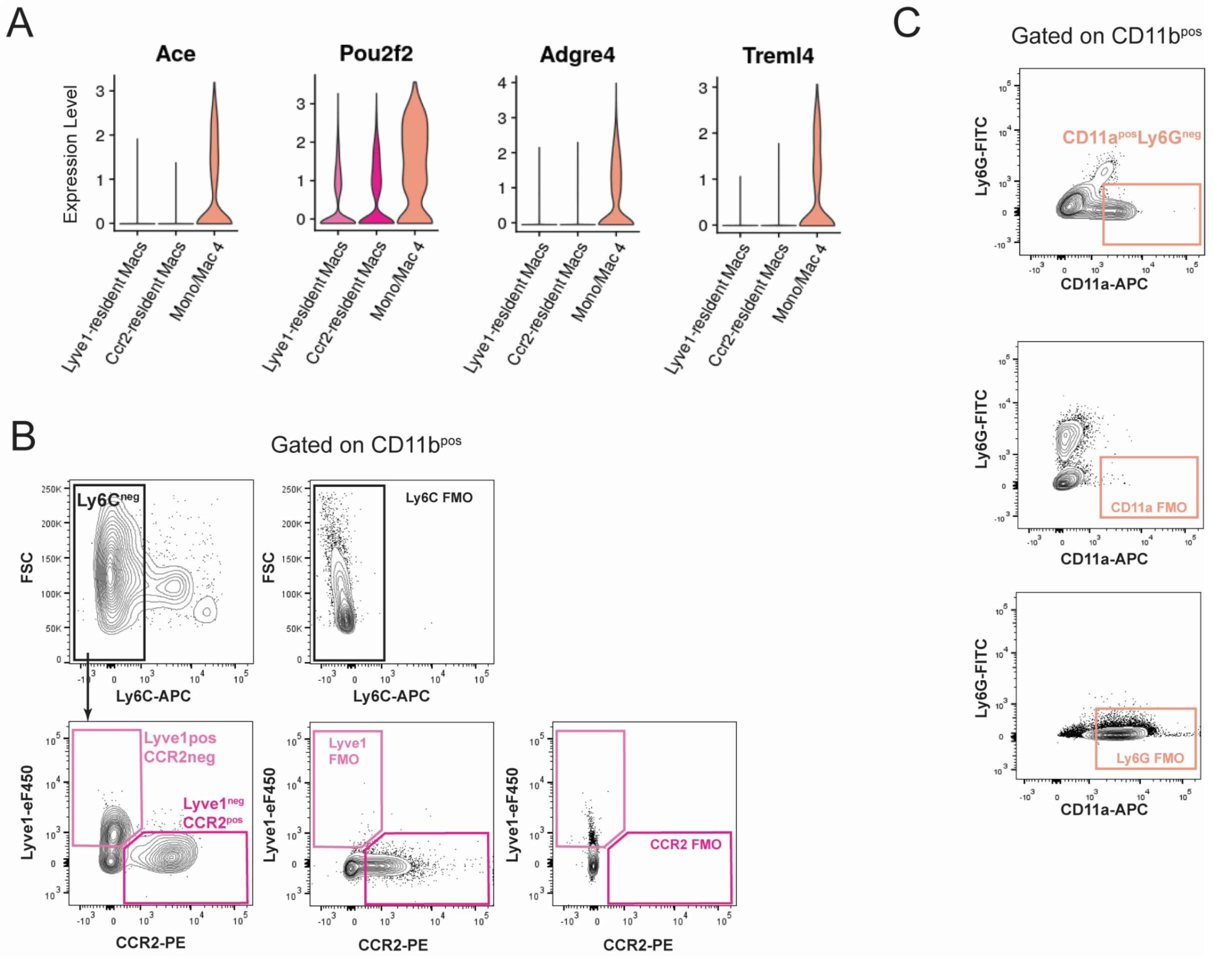


**Fig S6 | Macrophage populations in homeostatic skeletal muscle**

**A,** Violin plots showing normalized and log-transformed expression of representative markers of Mono/Mac 4 population across clusters of macrophages present in n.i. SkM of yg and old mice.

**B,** Gating strategy used in flow cytometry analysis of resident macrophage populations in n.i. SkM of yg and old mice, gated on CD11b$^{pos}$ cells: Lyve1-resident macrophages (Ly6C$^{neg}$Ccr2$^{neg}$Lyve1$^{pos}$) and Ccr2-resident macrophages (Ly6C$^{neg}$Lyve1$^{neg}$Ccr2$^{pos}$). The FMO density plots displayed were used to define the positive populations.

**C,** Gating strategy used in flow cytometry analysis of Mono/Mac4 population (Ly6G$^{neg}$CD11a$^{pos}$) in n.i. SkM of yg and old mice, gated on CD11b$^{pos}$ cells: Mono/Mac4 (Ly6G$^{neg}$CD11a$^{pos}$). The FMO density plots displayed were used to define the positive populations.

SkM, skeletal muscle; yg, young; n.i., non-injured; FSC, Forward Scatter; FMO, Fluorescence Minus One Control.

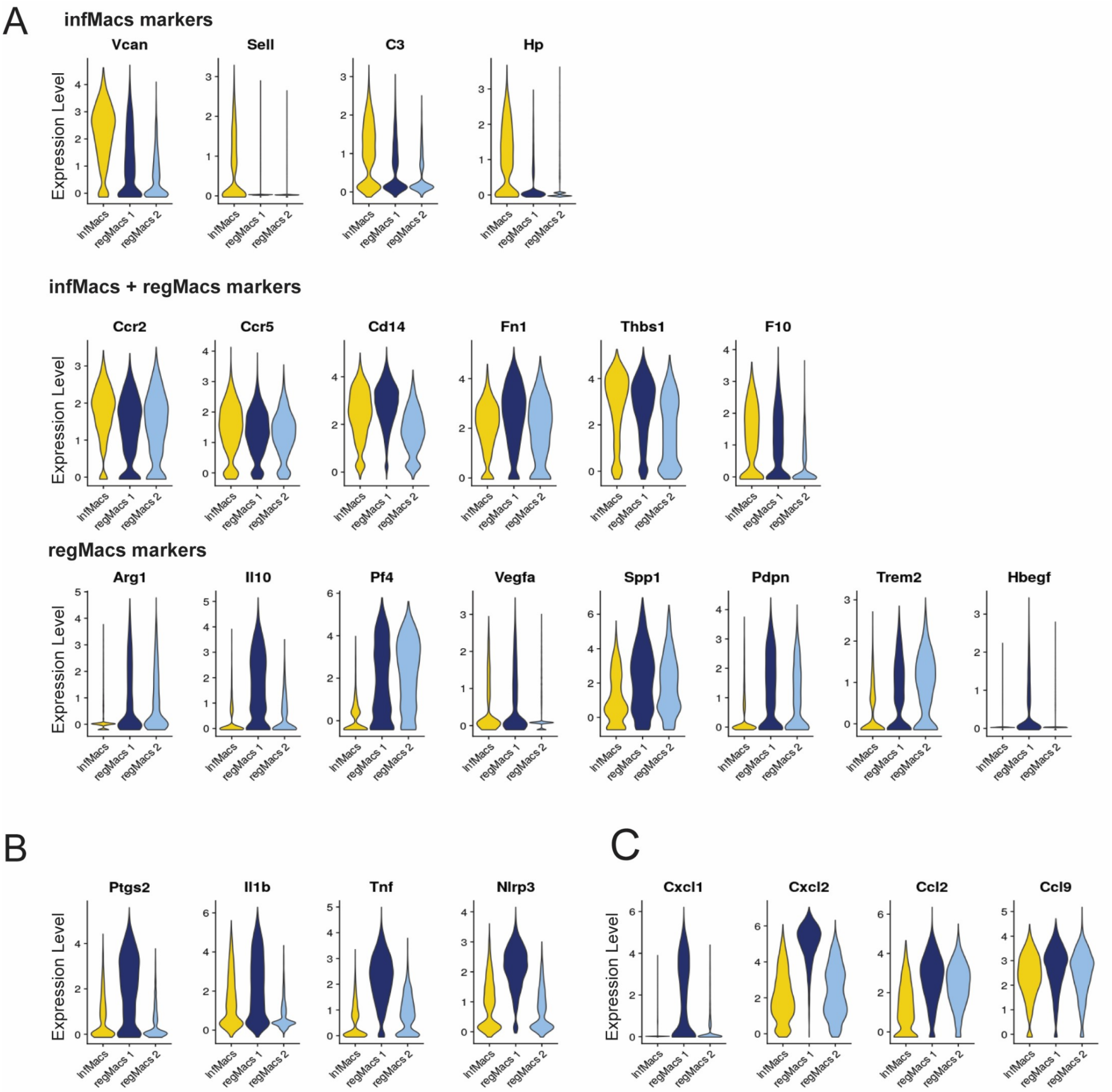


**Fig. S7 | Distributions of marker gene expression levels of macrophages during early regeneration.**

**A,** Violin plots showing normalized and log-transformed expression of representative markers of infMacs, regMacs 1 and regMacs 2, across all three clusters of macrophages detected in SkM of yg and old mice at 1dpi, as shown in Fig. 5B.

**B,** Violin plots showing normalized and log-transformed expression of pro-inflammatory markers across all three clusters of macrophages detected in SkM of yg and old mice at 1dpi, as shown in Fig. 5B.

**C,** Violin plots showing normalized and log-transformed expression of chemokines across all three clusters of macrophages detected in SkM of yg and old mice at 1dpi, as shown in Fig. 5B.

SkM, skeletal muscle; yg, young; dpi, days post injury; infMacs, infiltrating monocyte-derived macrophages; regMacs, regeneration-associated regulatory macrophages.

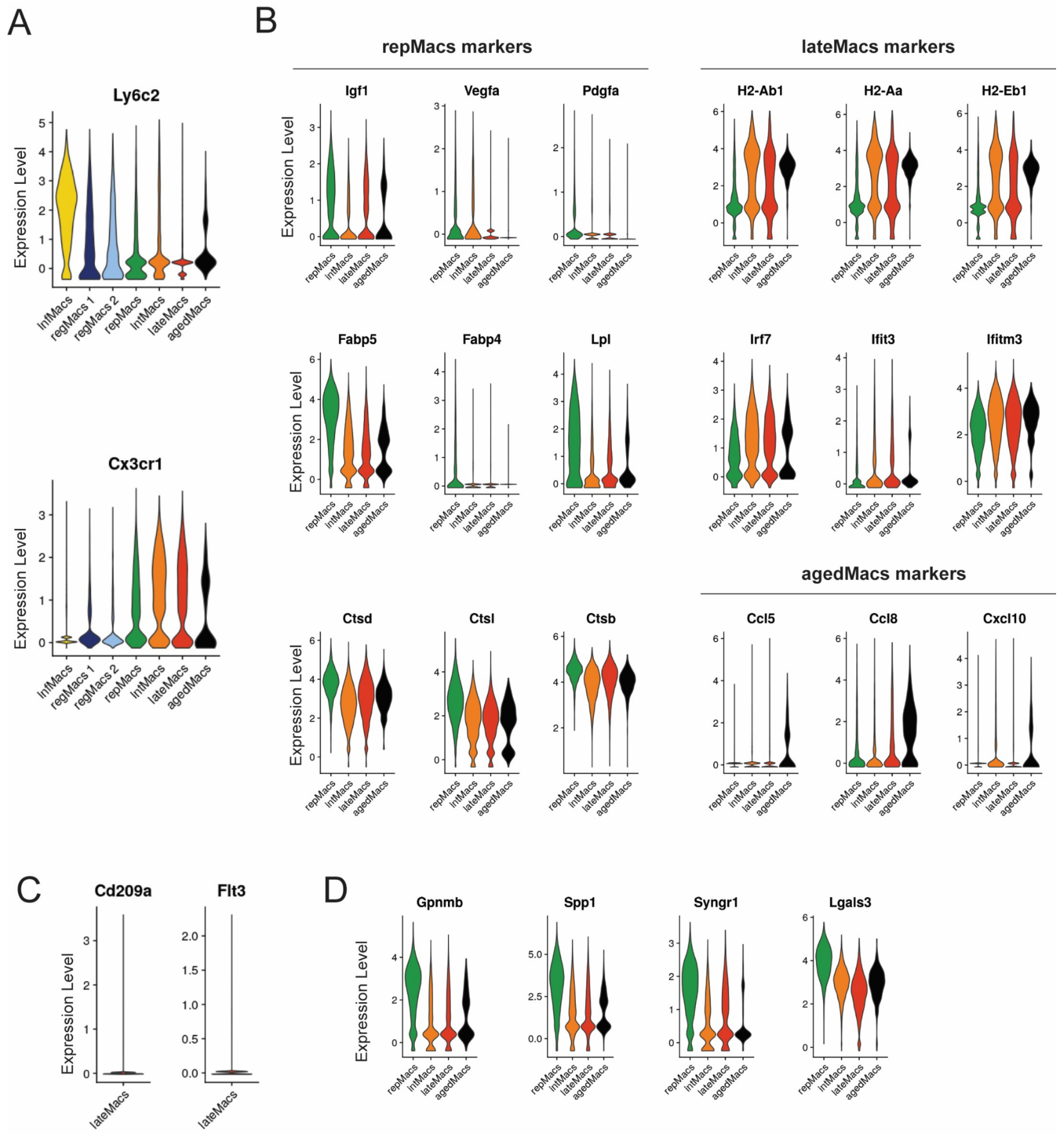


**Fig. S8 | Distributions of marker gene expression levels of macrophages during late regeneration.**

**A,** Violin plots showing normalized and log-transformed expression of Ly6c2 and Cx3cr1, across all clusters of macrophages detected in regenerating SkM of yg and old mice, as shown in Fig. 5B.

**B,** Violin plots showing normalized and log-transformed expression of representative markers of repMacs, lateMacs and ageMacs, across all four clusters of macrophages detected in SkM of yg and old mice at 3 and 5 dpi, as shown in Fig. 5B.

**C,** Violin plots showing normalized and log-transformed expression of DC markers in lateMacs.

**D,** Violin plots showing normalized and log-transformed expression of pro-fibrogenic markers, across all clusters of macrophages detected in regenerating SkM of yg and old mice, as shown in Fig. 5B.

SkM, skeletal muscle; yg, young; dpi, days post injury; infMacs, infiltrating monocyte-derived macrophages; regMacs, regeneration-associated regulatory macrophages; repMacs, repair-associated macrophages; intMacs, intermediate macrophages; lateMacs late regeneration-associated macrophages; ageMacs, age-associated macrophages; DCs, dendritic cells.

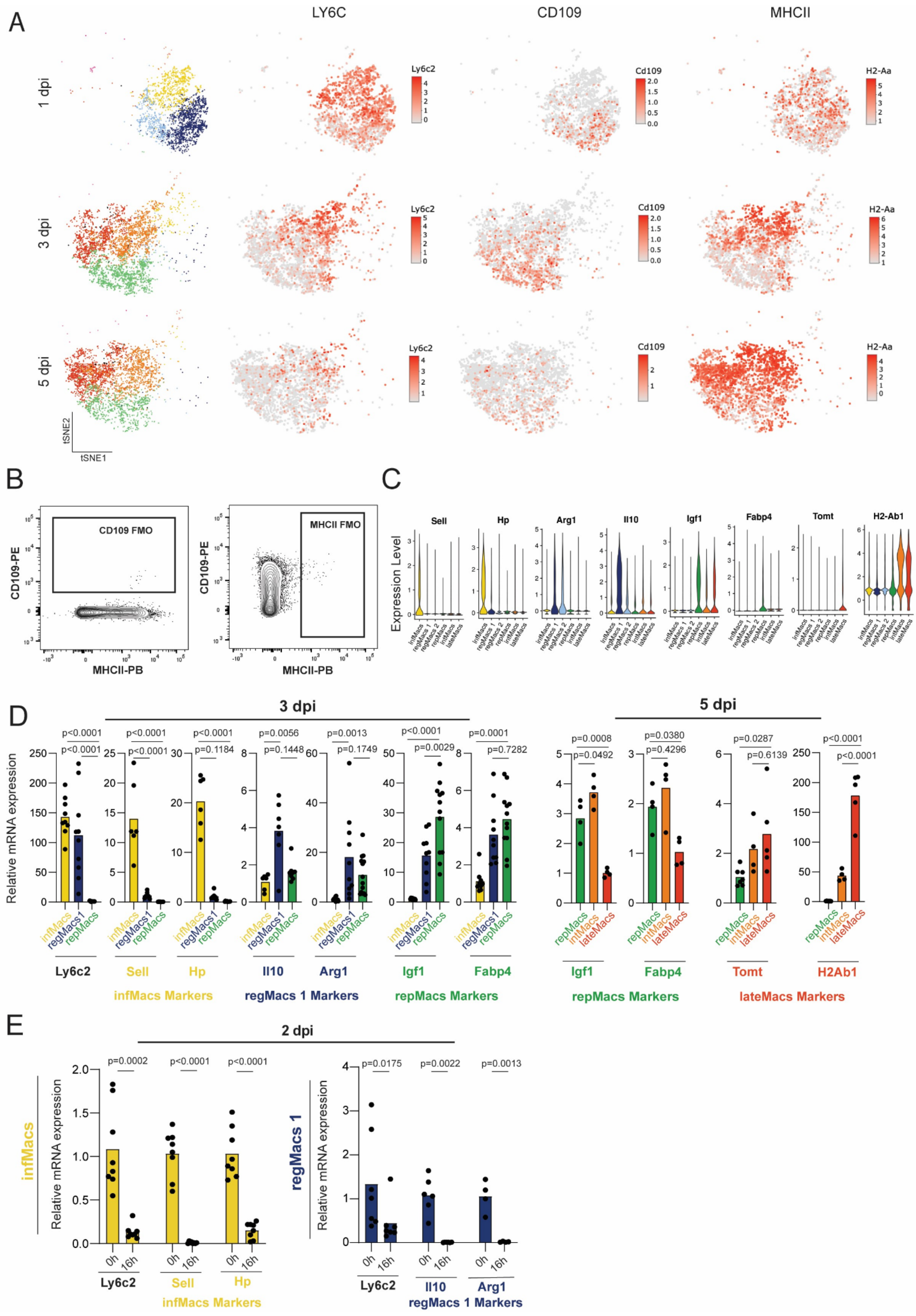
A
LY6C
CD109
MHCII
1 dpi
3 dpi
5 dpi
Ly6c2
Cd109
H2-Aa
tSNE2
tSNE1
B
CD109 FMO
MHCII FMO
CD109-PE
MHCII-PB
C
Expression Level
Sell
Hp
Arg1
Il10
Igf1
Fabp4
Tomt
H2-Ab1
infMacs
regMacs 1
regMacs 2
repMacs
intMacs
lateMacs
D
3 dpi
5 dpi
Relative mRNA expression
Ly6c2
Sell
Hp
Il10
Arg1
Igf1
Fabp4
Tomt
H2Ab1
infMacs Markers
regMacs 1 Markers
repMacs Markers
lateMacs Markers
E
2 dpi
infMacs
regMacs 1
0h
16h
infMacs Markers
regMacs 1 Markers

**Fig. S9 | Macrophage Dynamics during skeletal muscle regeneration**

**A,** tSNE plot showing the normalized and log-transformed expression levels of genes of interest to discriminate macrophage populations in regenerating SkM of yg mice, as defined by the re-clustering analysis of macrophage populations detected in the scRNAseq analysis, as shown in Fig. 5A-B.

**B,** FMO density plots used to define the positive populations in the flow cytometry panel shown in Fig. 6A.

**C,** Violin plots showing normalized and log-transformed expression of population-specific markers across all clusters of macrophages detected in regenerating SkM of yg and old mice, as shown in Fig. 5B.

**D,** Relative levels of Ly6c2, Sell, Hp, Il10, Arg1, Igf1, Fabp4, Tomt and H2Ab1 mRNA, detected by RT-qPCR, in infMacs, regMacs1, repMacs, intMacs and lateMacs, isolated, by FACS, from 3 and 5 dpi SkMs of yg wt (C57BL/6) mice (For 3 dpi analysis: n=7 for Sell, Hp and Il10 analysis at regMacs1 analysis at 3dpi; n=6 for Sell, Hp and Il10 analysis at infMacs; n=8 for Sell, Hp and Il10 analysis at repMacs; n=10 for Ly6c2, Arg1, Igf1 and Fabp4 analysis at regMacs1; n=9 for Ly6c2, Igf1 and Fabp4 analysis at infMacs; n=12 for Ly6c2, Igf1 and Fabp4 analysis at repMacs; n=11 for Arg1 analysis at infMacs; n=14 for Arg1 analysis at repMacs; For 5 dpi analysis: n=7 for Tomt and H2Ab1 analysis at repMacs; n=5 for Tomt and H2Ab1 analysis at lateMacs; n =4 for all other analysis).

**E,** Relative levels of Ly6c2, Sell, Hp, Il10 and Arg1 mRNA, detected by RT-qPCR, in infMacs and regMacs isolated, by FACS, from 2 dpi SkMs of yg wt (C57BL/6) mice, after 0h and 16h of in vitro culture (n=8 for analysis in infMacs; n=7 for Ly6c2 analysis in regMacs; n=6 for Il10 analysis in regMacs1; n=4 for Arg1 analysis in regMacs1).

In D and E data are represented as average, and each dot represents one animal.

In D, for Hp, Il10 and Fabp4 analysis at 3dpi, p-values are from Kruskal-Wallis test with Dunn´s post test. For all the other analyses, p values are from one-way ANOVA with Tukey post test. In E, for Ly6c2 analysis in infMacs and regMacs and for Il10 analysis in regMacs, p values are from two-tailed Mann-Whitney test. For all other analyses, p-values are from two-tailed Student’s t test.

SkM, skeletal muscle; yg, young; dpi, days post injury; infMacs, infiltrating monocyte-derived macrophages; regMacs, regeneration-associated regulatory macrophages; repMacs, repair-associated macrophages; intMacs, intermediate macrophages; lateMacs late regeneration-associated macrophages; ageMacs, age-associated macrophages; FMO, Fluorescence Minus One Control; MHC II, major histocompatibility complex class II.

A

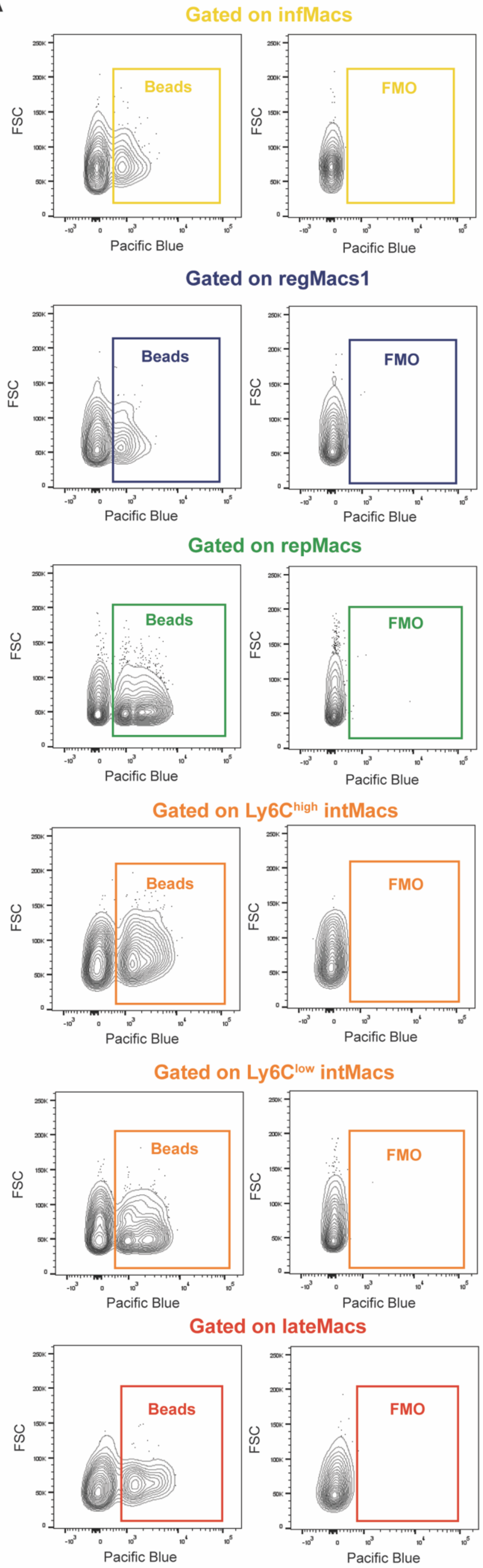
Gated on infMacs
Beads
FMO
FSC
Pacific Blue
Gated on regMacs1
Beads
FMO
FSC
Pacific Blue
Gated on repMacs
Beads
FMO
FSC
Pacific Blue
Gated on Ly6Chigh intMacs
Beads
FMO
FSC
Pacific Blue
Gated on Ly6Clow intMacs
Beads
FMO
FSC
Pacific Blue
Gated on lateMacs
Beads
FMO
FSC
Pacific Blue

**Fig. S10 | Flow cytometry analysis of phagocytosis by macrophages**

**A,** Gating strategy used in the flow cytometry analysis of Bright Blue-conjugated opsonized-beads ingestion by different macrophage populations, defined by the gating strategy shown in Fig. 6A. The FMO density plots displayed were used to define the phagocytic fraction for each macrophage subtype.

FSC, Forward Scatter; FMO, Fluorescence Minus One Control.

A

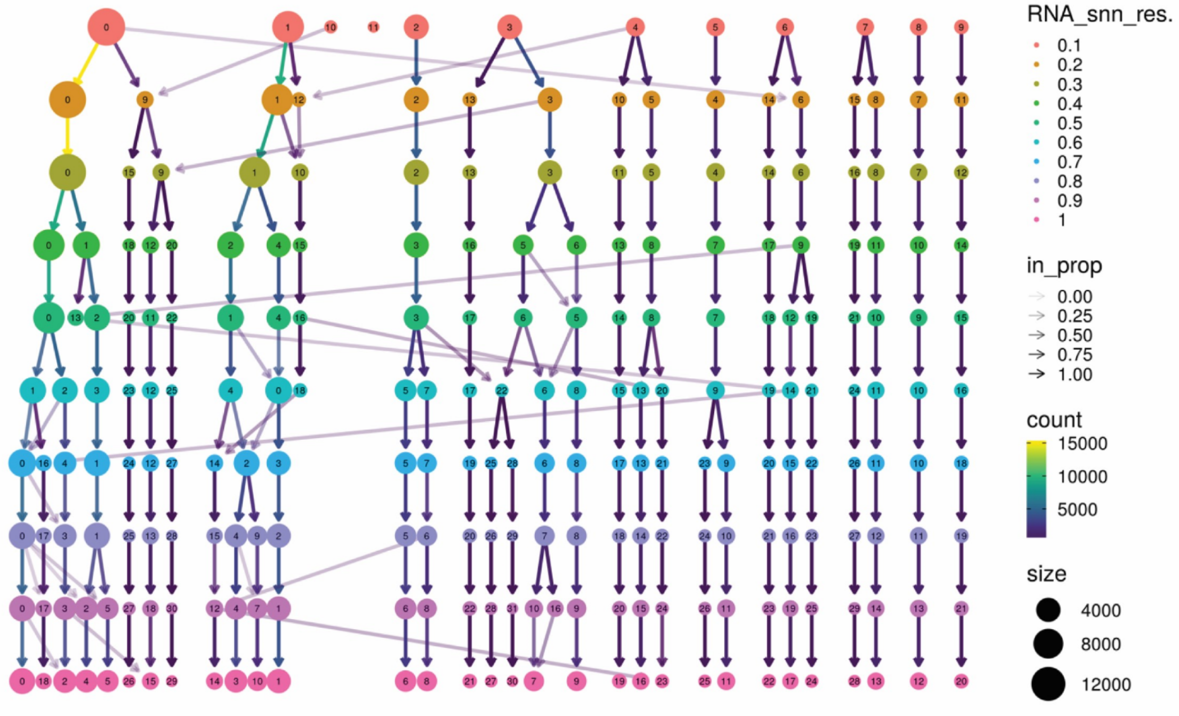
RNA_snn_res.
0.1
0.2
0.3
0.4
0.5
0.6
0.7
0.8
0.9
1
in_prop
0.00
0.25
0.50
0.75
1.00
count
15000
10000
5000
size
4000
8000
12000

B

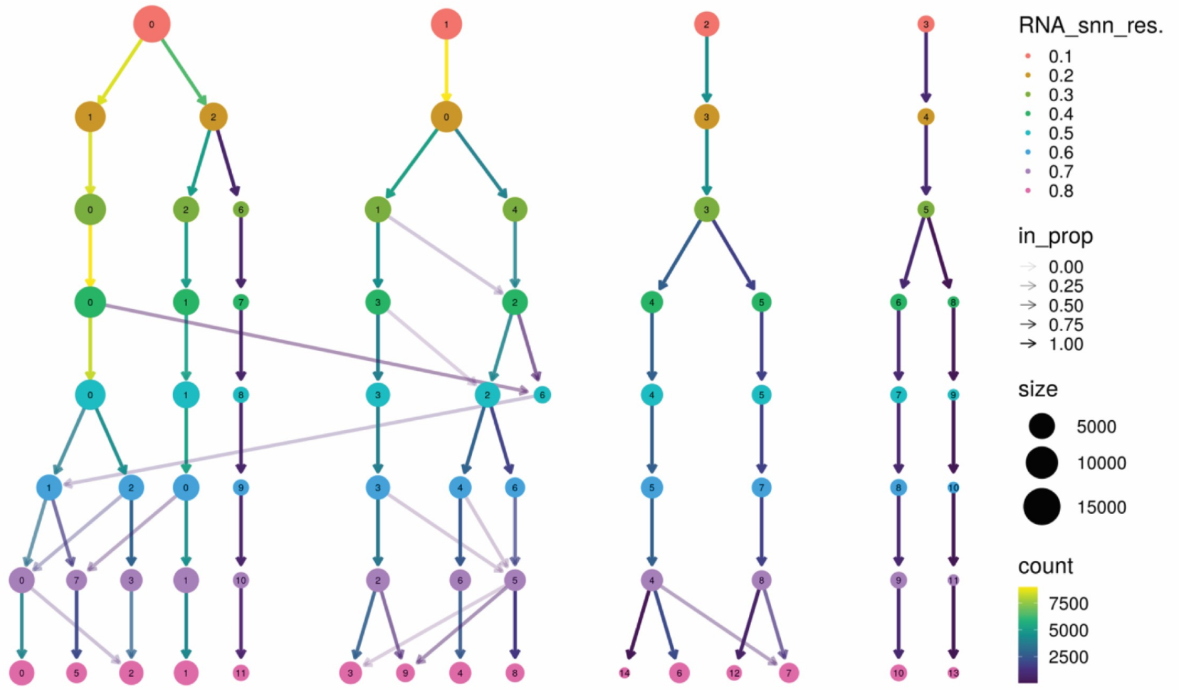
RNA_snn_res.
0.1
0.2
0.3
0.4
0.5
0.6
0.7
0.8
in_prop
0.00
0.25
0.50
0.75
1.00
size
5000
10000
15000
count
7500
5000
2500

**Fig. S11 | Cluster trees for scRNAseq analysis**

**A,** Cluster tree for the dataset comprising all cells.

**B,** Cluster tree for the dataset comprising only macrophages.

In this representation, every cluster forms a node in the tree, and edges between nodes are established by taking into account cells in a cluster at a lower resolution that transition to a cluster at the next higher resolution. The visualization parameters encompass the following attributes: node color corresponds to clustering resolution, edge opacity reflects the proportion of cells transitioning to the child cluster, edge color reflects the number of cells transitioning to the child cluster, node size represents the number of cells within the cluster.

**Table S1**

Fluorophore-conjugated antibodies for FC analysis and FACS

| Fluorophore | Antibody | Source | Dilution |
|---|---|---|---|
| **APC** | Anti-mouse CD197 (CCR7) | Biolegend (120107) | 1:20 |
| **APC** | Anti-mouse CD11a | Biolegend (141010) | 1:100 |
| **APC** | Anti-mouse SiglecF | Biolegend (155508) | 1:80 |
| **APC** | Anti-mouse CD209a | Biolegend (833005) | 1:20 |
| **APC** | Anti-mouse NK1.1 | Biolegend (108710) | 1:25 |
| **APC-eFluor®780** | Anti-mouse CD11b | eBioscience™ (47-0112-82) | 1:200 |
| **Brilliant Violet 421™** | Anti-mouse SiglecF | BD Horizon™ (562681) | 1:200 |
| **Brilliant Violet 711** | Anti-mouse LY6C | Biolegend (128037) | 1:40 |
| **Brilliant Violet 711** | Anti-mouse I-A/I-E | Biolegend (107643) | 1:80 |
| **Brilliant Violet 785™** | Anti-mouse CD3ε | Biolegend (100355) | 1:50 |
| **eF450™** | Anti-mouse LYVE1 | eBioscience™ (48-0443-82) | 1:20 |
| **FITC** | Anti-mouse CD79b | Biolegend (132805) | 1:200 |
| **FITC** | Anti-mouse XCR1 | Biolegend (148209) | 1:50 |
| **FITC** | Anti-mouse LY6G | Biolegend (127606) | 1:400 |
| **FITC** | Anti-mouse LY6C | Biolegend (128006) | 1:200 |
| **Pacific Blue ™** | Anti-mouse CD103 | Biolegend (121417) | 1:50 |
| **Pacific Blue ™** | Anti-mouse I-A/I-E | Biolegend (107620) | 1:200 |
| **PE** | Anti-mouse ST2 | Biolegend (145303) | 1:20 |
| **PE** | Anti-mouse CD209a | Biolegend (833004) | 1:20 |
| **PE** | Anti-mouse CXCR2 | Biolegend (149304) | 1:20 |
| **PE** | Anti-mouse CD109 | Bio-Techne (FAB4385P) | 1:10 |
| **PE** | Anti-mouse CCR2 | Biolegend (150610) | 1:40 |
| **PE-CY5** | Anti-mouse CD45 | Biolegend (103109) | 1:100 |
| **PE-Cy7** | Anti-mouse LY6G | Biolegend (127618) | 1:100 |

**Table S2**

Primer list for RT-qPCR

| Gene | Species | Forward Primer | Reverse Primer |
|---|---|---|---|
| **Beta-actin** | mouse | GCTCTGGCTCCTAGCACCAT | GCCACCGATCCACACAGAGT |
| **CD7** | mouse | CTTTGCTGCTTACACTGGC | TCAGAGGCAATCGTGAGTC |
| **Mrc1** | mouse | TTGTGGAGCAGATGGAAGGT | GTACATGGCTTCATATCCTCTCG |
| **Dcstamp** | mouse | TCTCCTCCATGAACAAACAG | CTTGGGTTCCTTGCTTCTC |
| **Ly6C2** | mouse | AGTACTCACGCTACAAAGTC | CATAGCACTCGTAGCACTG |
| **Sell** | mouse | CTTGGGACTGCAGAAACAC | ACTTCTGTTTGTCTCTTGGC |
| **Hp** | mouse | CTGTTGTCACTCTCCTGCT | AGCTGTCATCTTCAAAGTCCA |
| **Il10** | mouse | GTGGAGCAGGTGAAGAGTGA | CGAGGTTTTCCAAGGAGTTG |
| **Arg1** | mouse | GAACTGAAAGGAAAGTTCCCA | AATGTACACGATGTCTTTGGC |
| **Igf1** | mouse | TTTACTTCAACAAGCCCACAG | GGAAGCAACACTCATCCAC |
| **Fabp4** | mouse | CAGAAGTGGGATGGAAAGTC | GCCTTTCATAACACATTCCAC |
| **Tomt** | mouse | CCCTGTTAAAGGTCAGATTCTG | ATCCACAGTATGTGCCCAG |
| **H2Ab1** | mouse | CATCACTGTGGAGTGGAGG | TTTCTGACTCCTGTGACGG |
